# Structural Basis for Cytoplasmic Dynein-1 Regulation by Lis1

**DOI:** 10.1101/2021.06.11.448119

**Authors:** John P. Gillies, Janice M. Reimer, Eva P. Karasmanis, Indrajit Lahiri, Zaw Min Htet, Andres E. Leschziner, Samara L. Reck-Peterson

## Abstract

The lissencephaly 1 gene, *LIS1*, is mutated in patients with the neurodevelopmental disease lissencephaly. The Lis1 protein is conserved from fungi to mammals and is a key regulator of cytoplasmic dynein-1, the major minus-end-directed microtubule motor in many eukaryotes. Lis1 is the only dynein regulator that binds directly to dynein’s motor domain, and by doing so alters dynein’s mechanochemistry. Lis1 is required for the formation of fully active dynein complexes, which also contain essential cofactors: dynactin and an activating adaptor. Here, we report the first high-resolution structure of the yeast dynein–Lis1 complex. Our 3.1Å structure reveals, in molecular detail, the major contacts between dynein and Lis1 and between Lis1’s ß-propellers. Structure-guided mutations in Lis1 and dynein show that these contacts are required for Lis1’s ability to form fully active human dynein complexes and to regulate yeast dynein’s mechanochemistry and in vivo function. We present a model for the conserved role of Lis1 in regulating dynein from yeast to humans.

## Introduction

Cytoplasmic dynein-1 (dynein) is a minus-end-directed microtubule-based motor that is conserved across eukaryotes. Mutations in dynein or its regulators cause a range of neurodevelopmental and neurodegenerative diseases in humans (Lipka et al., 2013). In metazoans and some fungi, dynein transports organelles, RNAs, and protein complexes (Reck-Peterson et al., 2018). Dynein also positions nuclei and centrosomes and has a role in organizing the mitotic spindle and in the spindle assembly checkpoint (Raaijmakers and Medema, 2014). Given the essential nature of dynein in many organisms, the yeast *S. cerevisiae* has been an important model system for studying dynein’s mechanism and function, as dynein and its regulators are conserved, but non-essential in this organism. In yeast, deletion of dynein or dynein’s regulators results in defects in positioning the mitotic spindle (Eshel et al., 1993; Kormanec et al., 1991; Lee et al., 2003; Li et al., 2005, 1993; Muhua et al., 1994; Sheeman et al., 2003).

Both dynein and its regulation are complex. Cytoplasmic dynein is a dimer of two motor-containing AAA+ (<u>A</u>TPase <u>a</u>ssociated with various cellular <u>a</u>ctivities) subunits, with additional subunits all present in two copies. In the absence of regulatory factors dynein exists largely in an autoinhibited “phi” conformation (Torisawa et al., 2014; Zhang et al., 2017). Active moving dynein complexes contain the dynactin complex and a coiled-coil-containing activating adaptor (Mckenney et al., 2014; Schlager et al., 2014; Trokter et al., 2012) (Figure 1A). Some of these activated dynein complexes contain two dynein dimers, which move at a faster velocity than complexes containing a single dynein dimer (Elshenawy et al., 2020; Grotjahn et al., 2018; Htet et al., 2020; Urnavicius et al., 2018). About a dozen activating adaptors have been described in mammals, including members of the Hook, Bicaudal D (BicD), and ninein families (Canty and Yildiz, 2020; Olenick and Holzbaur, 2019; Reck-Peterson et al., 2018). In yeast, the coiled-coil-containing protein Num1 is required to activate dynein in vivo (Heil-Chapdelaine et al., 2000; Lammers and Markus, 2015; Lee et al., 2003; Sheeman et al., 2003) and is likely a dynein activating adaptor.

**Figure 1.**
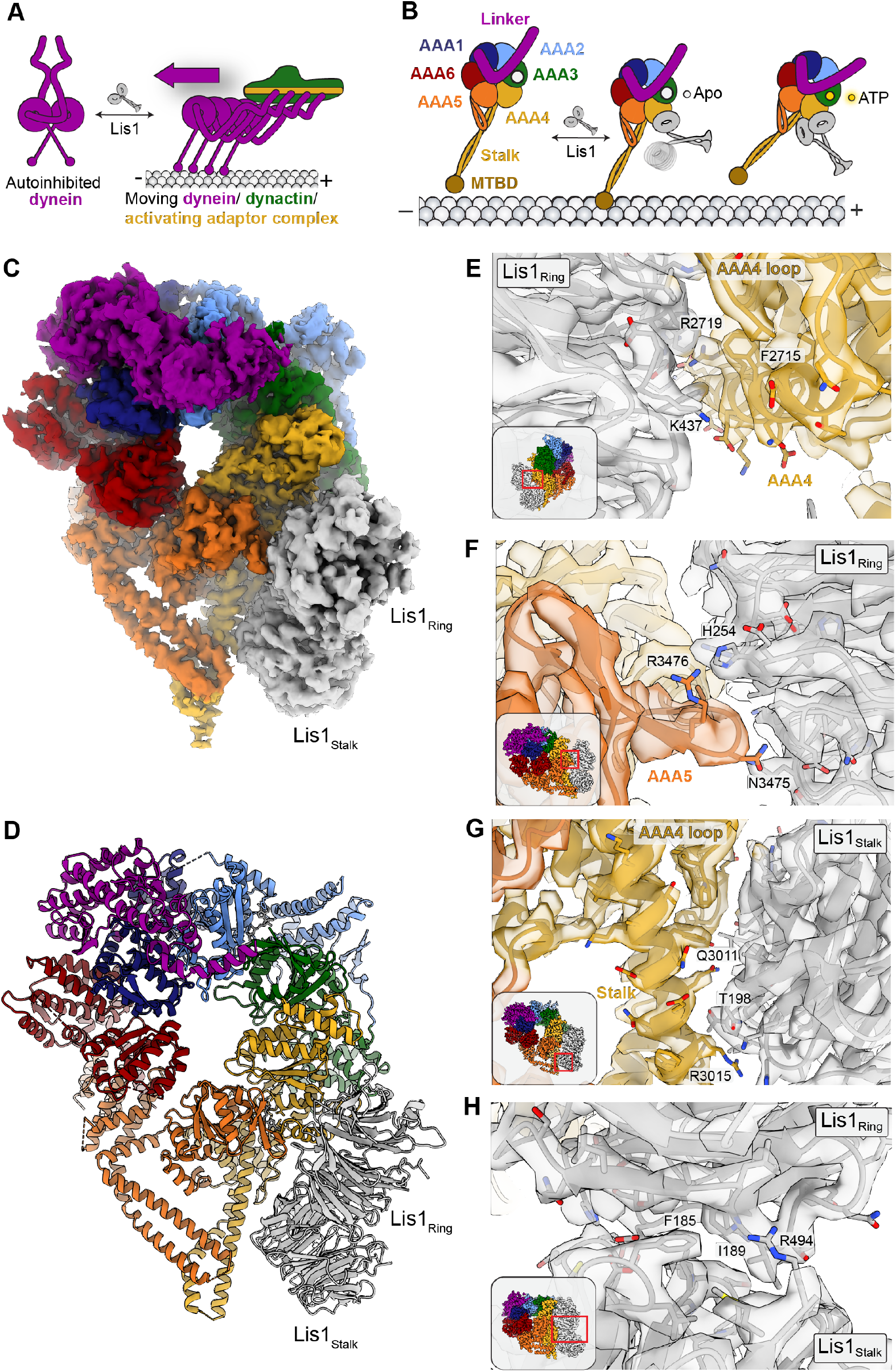
A 3.1Å structure of the dynein-Lis1 complex. **(A)** Lis1 assists in the transition from autoinhibited dynein to activated dynein complexed with dynactin and an activating adaptor protein. **(B)** Schematic representation of the different modes of binding of Lis1 to dynein as a function of nucleotide state at AAA3. **(C, D)** Cryo-EM map (C) and model (D) of dynein AAA3-WalkerB mutant with Lis1 bound at site_ring_ and site_stalk_. **(E-H)** Close-up views of the interfaces between dynein and Lis1 and of the Lis1_ring_-Lis1_stalk_ interface. Key residues are indicated for the site_ring_ interaction at AAA4 (E), the site_ring_ interaction at AAA5 (F), the site_stalk_ interaction (G) and the Lis1_ring_-Lis1_stalk_ interface (H), with insets showing the location of the close-up in the full structure.

In addition to dynactin and an activating adaptor, dynein function in vivo also requires Lis1. Genetically, Lis1 is a positive regulator of dynein function (Geiser et al., 1997; Liu et al., 1999; Xiang et al., 1995) and is required for most, if not all, dynein functions in many organisms (Markus et al., 2020). In humans, mutations in both Lis1 (*PAFAH1B1*) and the dynein heavy chain (*DYNC1H1*) cause the neurodevelopmental disease lissencephaly (Parrini et al., 2016; Reiner et al., 1993). Structurally, Lis1 is a dimer of two ß-propellers (Kim et al., 2004; Tarricone et al., 2004) and is the only known dynein regulator that binds directly to dynein’s motor domain (Huang et al., 2012; Toropova et al., 2014). Structures of dynein–Lis1 complexes, while not of high enough resolution to build molecular models, show that Lis1 binds to dynein at two sites: dynein’s AAA+ ring at AAA3/AAA4 (site_ring_) and dynein’s stalk (site_stalk_), which emerges from AAA4 and leads to dynein’s microtubule binding domain (Figure 1B) (DeSantis et al., 2017; Htet et al., 2020; Huang et al., 2012; Toropova et al., 2014).

Given that Lis1 directly binds to two different regions of the dynein motor, how does Lis1 impact dynein motility, and ultimately function as a positive regulator of dynein motility? Recently, Lis1 was shown to play a role in forming active dynein complexes (Elshenawy et al., 2020; Htet et al., 2020; Marzo et al., 2020; Qiu et al., 2019). Human dynein–dynactin–activating adaptor complexes formed in the presence of Lis1 are more likely to contain two dynein dimers and move at a faster velocity in vitro (Elshenawy et al., 2020; Htet et al., 2020). Studies in yeast and the filamentous fungus *Aspergillus nidulans* suggest that this role for Lis1 in forming active complexes is conserved and required for in vivo function (Marzo et al., 2020; Qiu et al., 2019). Prior to this series of discoveries, we and others had explored Lis1’s effects on dynein’s mechanochemistry in vitro. A common theme in these studies was that the presence of Lis1 causes dynein to bind microtubules more tightly under some nucleotide conditions (DeSantis et al., 2017; Htet et al., 2020; Huang et al., 2012; Mckenney et al., 2010; Yamada et al., 2008), which in the context of a moving molecule correlates with slowed velocity (DeSantis et al., 2017; Huang et al., 2012; Yamada et al., 2008) or increased stalling under load (Mckenney et al., 2010). The nucleotide state of dynein’s AAA3 seems to regulate this, as Lis1 induces a weak microtubule binding state in yeast dynein when the nucleotide bound at AAA3 is non-hydrolysable (DeSantis et al., 2017). These two modes of dynein regulation by Lis1 appear to be related to how Lis1 interacts with dynein, with one ß-propeller bound at site_ring_ in the absence of nucleotide at AAA3, and two ß-propellers bound at site_ring_ and site_stalk_ when the nucleotide at AAA3 is non-hydrolysable (DeSantis et al., 2017).

Despite this progress, a full understanding of how Lis1 promotes dynein complex assembly and how this is related to Lis1’s role in regulating dynein’s mechanochemistry has been hampered by the lack of high-resolution structures of dynein–Lis1 complexes. Here, we report a 3.1Å structure of yeast dynein bound to two Lis1 ß-propellers. This is the first high-resolution structure of the dynein–Lis1 complex and reveals in molecular detail the contacts between dynein and Lis1 at both site_ring_ and site_stalk_ and the interface between two Lis1 ß-propellers. We use this structure to interrogate the dynein–Lis1 interaction both in vitro and in vivo, showing that these contacts are important for dynein’s role in nuclear positioning in yeast and dynein’s mechanochemistry in vitro. Finally, we show that mutating site_ring_ and site_stalk_ in human dynein disrupts Lis1’s ability to promote the formation of activated human dynein complexes, demonstrating that the Lis1 binding sites on dynein that alter its mechanochemistry are the same sites that are important for dynein complex formation. Our results point to a conserved model for dynein regulation by Lis1.

## Results

### 3.1Å structure of a dynein-Lis1 complex

Previously, we determined structures of a *S. cerevisiae* dynein motor domain bound to one or two yeast Lis1 ß-propellers by cryo-EM at 7.7Å and 10.5Å resolution, respectively (DeSantis et al., 2017). Although those structures allowed us to dock homology models of Lis1 into the map and determine their general orientation, their limited resolutions prevented us from building molecular models into the density or rotationally aligning Lis1’s ß-propellers. In order to address these limitations and to understand the molecular features of the interactions between dynein and Lis1, we aimed to improve the overall resolution of the dynein–Lis1 complex. To enrich our sample for complexes containing two Lis1 ß-propellers bound to dynein, we used a construct (supplementary file 1) of the yeast dynein motor domain containing a Walker B mutation in AAA3 (E2488Q) that prevents the hydrolysis of ATP, as our previous work showed that this state results in Lis1 binding at both site_ring_ and site_stalk_ (DeSantis et al., 2017). As with our previous structural studies (DeSantis et al., 2017; Toropova et al., 2014), we used monomers of the dynein motor domain and Lis1 dimers, referred to as dynein–(Lis1)_2_ here. We also added ATP vanadate to our samples, as it is hydrolyzed to ADP-vanadate and mimics a post-hydrolysis state for dynein (Schmidt et al., 2015), referred to here as ADP-V_i_. Our initial attempts at obtaining a higher resolution structure were limited by dynein’s strong preferred orientation in open-hole cryo-EM grids. To overcome this, we randomly and sparsely biotinylated dynein and applied the sample to streptavidin affinity grids (Han et al., 2016, 2012; Lahiri et al., 2019); this tethering results in randomly oriented complexes that are prevented from reaching and interacting with the air-water interface.

This approach allowed us to determine the structure of dynein bound to Lis1 in the presence of ATP vanadate to a nominal resolution of 3.1Å (Figure 1C and D; Figure 1–figure supplement 1A–C; Movie 1; supplementary file 2). In our structure, Dynein’s AAA ring is “closed”, which is seen when dynein’s microtubule binding domain is low affinity, the linker is bent (Bhabha et al., 2014; Schmidt et al., 2015), and when two Lis1 ß-propellers interact with dynein (DeSantis et al., 2017). It is likely that the two ß-propellers come from the same homodimer as the aminotermini of the two ß-propellers are very close to each and we did not detect density for additional ß-propellers during data processing. This new high-resolution map allowed us to build an atomic model of the entire dynein–(Lis1)_2_ complex. Binding of Lis1 at site_ring_ primarily involves the smaller face of the ß-propeller in Lis1 and an alpha helix in AAA4 of dynein (Figure 1E). Our high-resolution structure also allowed us to unambiguously map an additional interaction between dynein and Lis1 at site_ring_ that involves a contact with a loop in AAA5 (Figure 1F). Lis1 binds to dynein at site_stalk_ using its side and makes interactions with coiled-coil 1 (CC1) of dynein’s stalk, as well as a novel contact with a loop in AAA4 (residues 2935-2942) (Figure 1G). As suggested by the continuous density in our previous low-resolution structure (DeSantis et al., 2017), the Lis1s bound at site_ring_ and site_stalk_ directly interact with each other; this interface involves van der Waals and hydrophobic contacts (Figure 1H). The ATP binding sites at AAA1, AAA2, AAA3, and AAA4 are all occupied by nucleotides (Figure 1–figure supplement 1D–G). While AAA4 is clearly bound to ADP (Figure 1–figure supplement 1G), we were unable to resolve whether ATP or ADP-vanadate was bound at AAA1-AAA3, and chose to model ATP into each binding pocket (Figure 1–figure supplement 1D– F).

Dynein is a dynamic protein and undergoes multiple conformational changes as part of its mechanochemical cycle. ATP binding and hydrolysis at AAA1 leads to a cascade of conformational changes including closing of the AAA ring, shifting of the buttress and stalk helices, and swinging of the linker domain during its power stroke cycle (Cianfrocco et al., 2015). While the AAA ring domain of dynein reached the highest resolution in our structure, the linker and stalk helices are at significantly lower resolutions, most likely due to conformational heterogeneity (Figure 1–figure supplement 1B). We used 3D variability analysis (Punjani and Fleet, 2021) in cryoSPARC to generate variability components that capture continuous movements within the complex; the first two components describe the major movements. The first variability component shows the linker pulling away from its docked site at AAA2 and starting to straighten towards AAA4 in what could be the beginning of the power stroke (Figure 1–figure supplement 2A, Movie 2). The second variability component shows the AAA ring “breathing” slightly between its open and closed conformations, with associated movements in the buttress and stalk helices (Figure 1–figure supplement 2B, Movie 2). Importantly, all the interactions involving Lis1—with site_ring_ and site_stalk_ and at the Lis1-Lis1 interface—are maintained in the different conformations.

### Lis1’s interaction with dynein’s AAA5 contributes to regulation at site_ring_

The bulk of the dynein-Lis1 interface at site_ring_ is comprised of a helix in AAA4 of dynein, mutations in which disrupt Lis1 binding to dynein (Huang et al., 2012). Here we set out to determine if the additional contact we observed at site_ring_ involving AAA5 (Figure 1F) also contributes to dynein regulation by Lis1. In order to disrupt this electrostatic interaction (Figure 2A) we either mutated Lis1’s Glu253 and His254 to Ala or deleted dynein’s Asn3475 and Arg3476, which are both present in a short loop (Figure 2A). These mutations were engineered into the endogenous copies of either Lis1 (*PAC1*) or dynein (*DYN1*). Western blots of strains containing epitope-tagged versions of the mutants confirmed that both Lis1^E253A, H254A^ and Dynein^ΔN3475, ΔR3476^ were expressed at wild type levels (Figure 2–figure supplement 1A–C). To determine if these mutations had a phenotype in vivo, we performed a nuclear segregation assay. In this assay, deletion of dynein or dynactin subunits or Lis1 causes an increase in binucleate mother cells, which arise from defects in mitotic spindle positioning (Eshel et al., 1993; Kormanec et al., 1991; Lee et al., 2003; Li et al., 1993; Muhua et al., 1994; Sheeman et al., 2003). We found that Lis1^E253A, H254A^ and dynein^ΔN3475, ΔR3476^ both exhibit a binucleate phenotype that is equivalent to that seen with deletion of Lis1 or dynein (Figure 2B–D), suggesting that the interaction between Lis1 and dynein at AAA5 is required for Lis1’s ability to regulate dynein in vivo.

**Figure 2.**
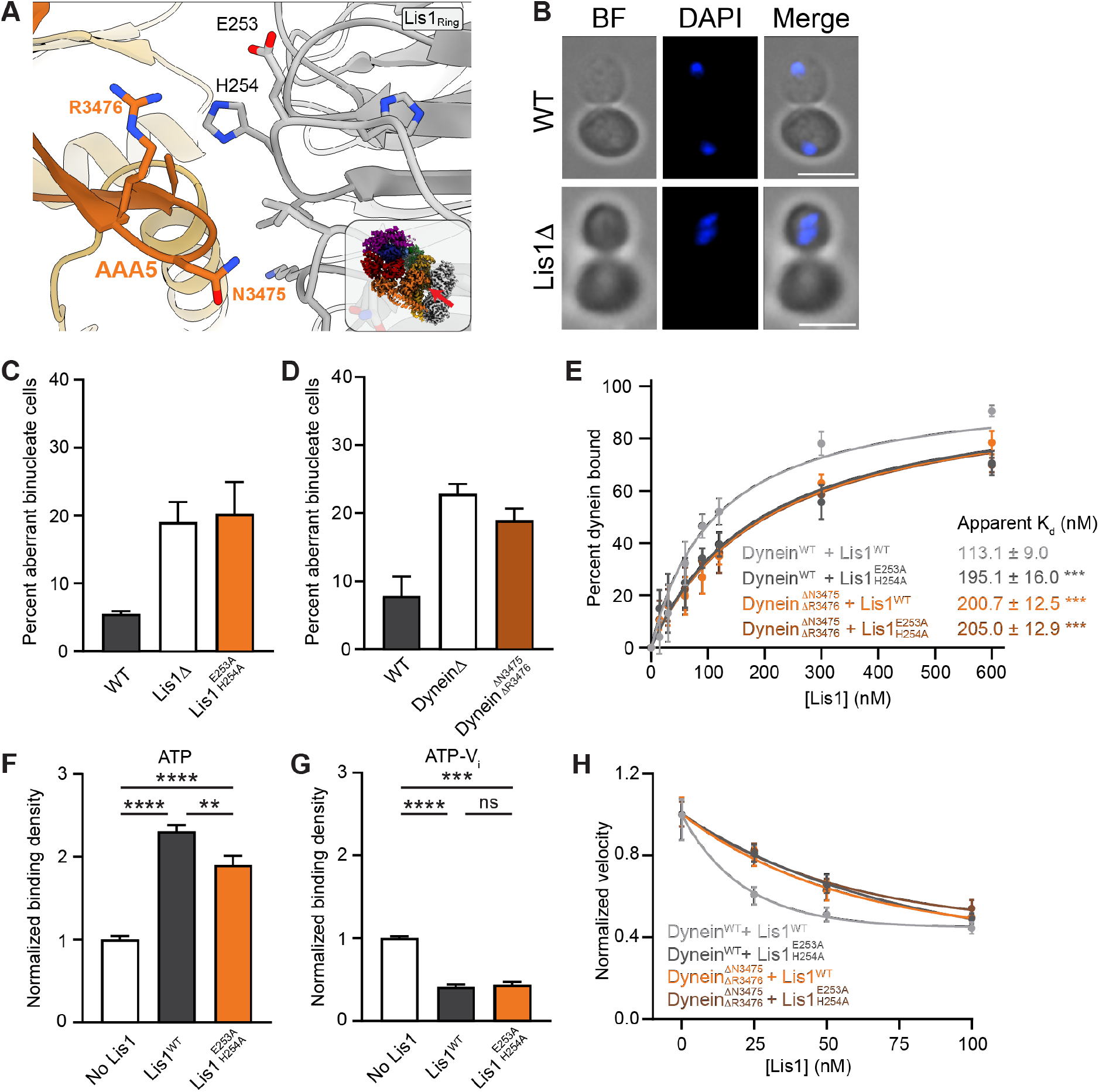
Lis1 regulation of dynein at site_ring_. **(A)** Close-up of the interface between Lis1 and dynein AAA5 at site_ring_. The residues mutated in this study on Lis1 (grey type) and dynein (orange type) are indicated. The inset shows the location of the close-up in the full structure. **(B)** Example images of normal and binucleate cells. Scale bar, 3μm **(C)** Quantitation (mean ± s.e.m.) of the percentage of cells displaying an aberrant binucleate phenotype for WT (dark grey), Lis1 deletion (white) and Lis1^E253A, H254A^ (orange). n = 3 replicates with at least 200 cells per condition. **(D)** Quantitation (mean ± s.e.m.) of the percentage of cells displaying an aberrant binucleate phenotype for WT (dark grey), Lis1 deletion (white) and dynein^ΔN3475, R3476^ (cark orange). n = 3 replicates with at least 200 cells per condition. **(E)** Quantitation of the apparent binding affinity of dynein^WT^ for Lis1^WT^ (dark grey; K_d_ = 113.1 ± 9.0) and Lis1^E253A, H254A^ (light gray; K_d_ = 195.1 ± 16.0) and dynein^ΔN3475, R3476^ for Lis1^WT^ (orange; K_d_ = 200.7 ± 12.5) and Lis1^E253A, H254A^ (brown; K_d_ = 205.0 ± 12.9). n = 3 replicates per condition. Statistical analysis was performed using an extra sum-of-squares F test; ***, p<0.0001. **(F)** Binding density (mean ± s.e.m.) of dynein on microtubules with ATP in the absence (dark grey) or presence of Lis1^WT^ (white) or Lis1^E253A, H254A^ (orange). Data was normalized to a density of 1.0 in the absence of Lis1. Statistical analysis was performed using an ANOVA; ****, p<0.0001; **, p = 0.0029. n = 12 replicates per condition. **(G)** Binding density (mean ± s.e.m.) of dynein on microtubules with ATP-V_i_ in the absence (dark grey) or presence of Lis1^WT^ (white) or Lis1^E253A, H254A^ (orange). Data was normalized to a density of 1.0 in the absence of Lis1. Statistical analysis was performed using an ANOVA; ****, p<0.0001; ***, p = 0.0001; ns = 0.9997. n = 12 replicates per condition. **(H)** Single-molecule velocity of dynein with increasing concentrations of Lis1. Dynein^WT^ with Lis1^WT^ (dark grey); dynein^WT^ with Lis1^E253A, H254A^ (light grey); dynein^ΔN3475, R3476^ with Lis1^WT^ (orange); dynein^ΔN3475, R3476^ and Lis1^E253A, H254A^ (brown). The median and interquartile range are shown. Data was normalized to a velocity of 1.0 in the absence of Lis1. At least 400 single molecule events were measured per condition.

We next characterized the biochemical properties of these mutations designed to disrupt Lis1’s interaction with AAA5. We began by measuring the apparent binding affinity between a Lis1 dimer and a monomeric dynein motor domain. For these experiments, purified SNAP-tagged Lis1 was covalently coupled to magnetic beads, mixed with purified dynein motor domains in the presence of ADP, and the percentage of dynein bound to the beads was quantified. We found that mutations at the Lis1-AAA5 interface, either on Lis1 (Lis1^E253A, H254A^) or on dynein (dynein^ΔN3475, ΔR3476^), led to ~45% decrease in Lis1 binding to dynein (Figure 2E). Next, we looked at Lis1’s effect on dynein’s interaction with microtubules. Previously, we showed that tight microtubule binding by dynein in the presence of ATP is mediated by site_ring_, whereas Lis1-induced weak microtubule binding in the presence of ATP-V_i_ requires both site_ring_ and site_stalk_ (DeSantis et al., 2017). For these experiments we used purified Lis1 dimers and purified truncated and GST-dimerized dynein motors (Reck-Peterson et al., 2006). These GST-dynein dimers (“dynein” here) have been well-characterized and behave similarly to full-length yeast dynein *in vitro* (Gennerich et al., 2007; Reck-Peterson et al., 2006). As expected for a mutation that would disrupt Lis1’s interaction at site_ring_, we found that Lis1^E253A, H254A^ decreased Lis1’s ability to enhance the binding of dynein to microtubules in the presence of ATP (Figure 2F). In contrast, Lis1^E253A, H254A^ did not affect Lis1’s ability to lower dynein’s affinity for microtubules in the presence of ATP-V_i_ (Figure 2G).

Finally, we measured the effect of disrupting the Lis1-AAA5 interaction on the velocity of dynein using a single-molecule motility assay. Unlike mammalian dynein, *S. cerevisiae* dynein moves processively in vitro in the absence of any other dynein subunits or cofactors (Reck-Peterson et al., 2006). We previously showed that addition of Lis1 to in vitro motility assays slows dynein in a dosedependent manner (Huang et al., 2012). Here, we found that in the absence of Lis1, dynein^ΔN3475, ΔR3476^ had a similar velocity to wild type dynein, indicating that the mutation does not affect the properties of the motor (Figure 2H). In contrast, Lis1 was less effective at slowing the velocity of dynein^ΔN3475, ΔR3476^ compared to wild type dynein. We observed a similar effect with Lis1^E253A, H254A^, which targets the same site_ring_ interface, but from the Lis1 side (Figure 2H). Thus, the interaction of Lis1 with dynein at AAA5 contributes to Lis1’s regulation of dynein at site_ring_.

### Lis1 regulation of dynein at site_ring_ does not require Lis1’s interaction with microtubules

A recent report proposed that yeast Lis1’s ability to slow dynein’s velocity might reflect a tethering mechanism, where Lis1 binds to both dynein and the microtubule (Marzo et al., 2020). This tethering model requires that either (1) one Lis1 β-propeller in a Lis1 dimer interact with dynein and the other with the microtubule (Figure 3A), or (2) a single Lis1 β-propeller interact simultaneously with both dynein and the microtubule (Figure 3B).

**Figure 3.**
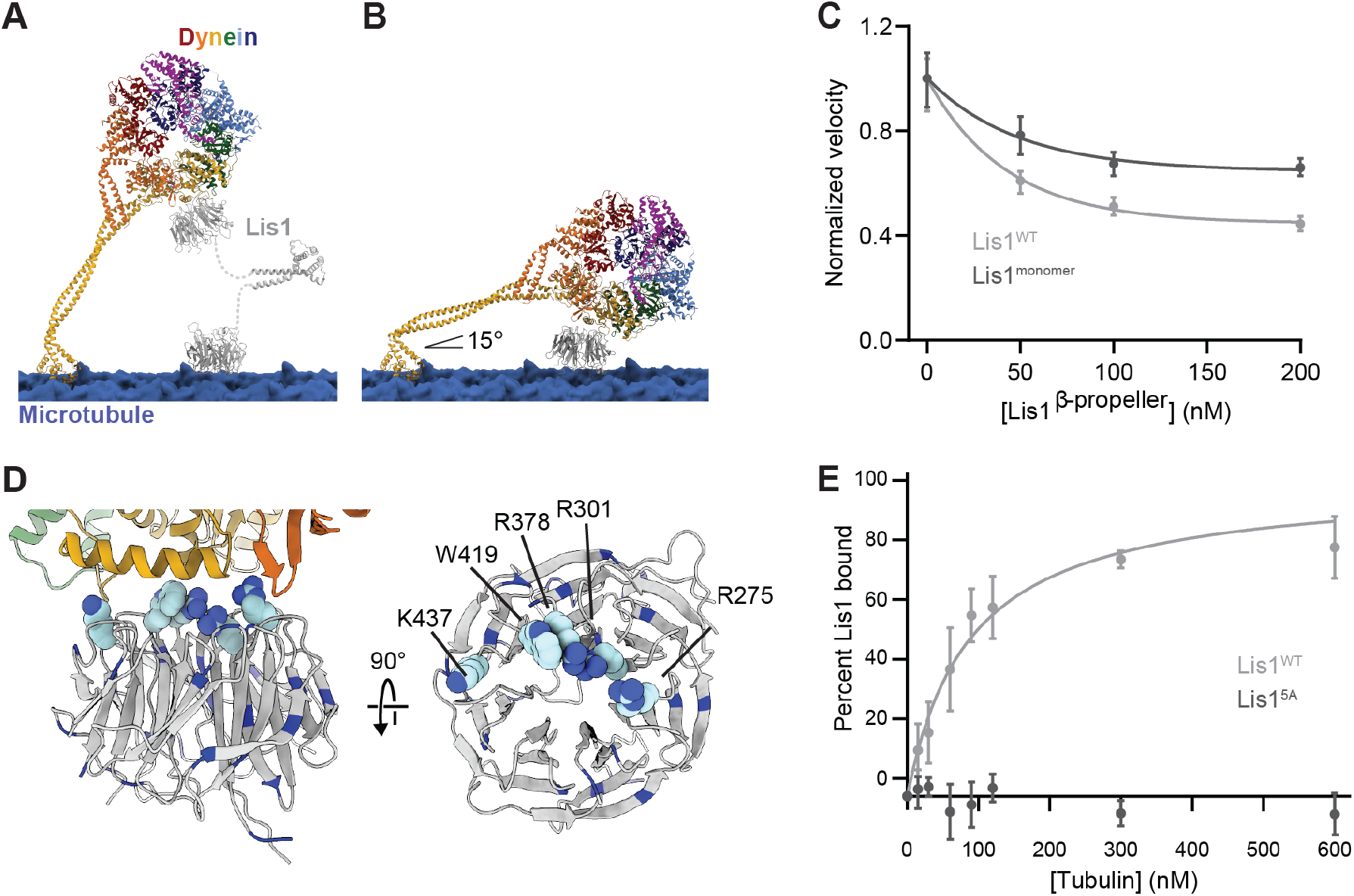
Lis1 regulation of dynein at site_ring_ does not require Lis1’s interaction with microtubules. **(A)** Model of human dynein 2 (PDB: 4RH7) (Schmidt et al., 2015, 2) tethered to a microtubule (blue) via a Lis1 dimer (grey), with one Lis1 ß-propeller docked to site_ring_ on dynein (PDB: 5VLJ) (DeSantis et al., 2017) and the other on the microtubule. **(B)** Model of human dynein 2 (PDB: 4RH7) (Schmidt et al., 2015, 2) with one Lis1 ß-propeller docked to site_ring_ on dynein (PDB: 5VLJ) (DeSantis et al., 2017) was docked into a tomogram of dynein on the microtubule (Grotjahn et al., 2018). In order to bring the Lis1 ß-propeller in contact with the microtubule, dynein’s stalk was bent near the two prolines found next to its microtubule binding domain. This results in an angle for the stalk of around 15°. **(C)** Single-molecule velocity of dynein with increasing concentrations of Lis1^WT^ (light grey) or Lis1^monomer^ (dark grey). The median and interquartile range are shown. Data was normalized to a velocity of 1.0 in the absence of Lis1. At least 400 single molecule events were measured per condition. **(D)** The structure of Lis1 at site_ring_ with the five residues mutated in Lis1^5A^ shown as spheres and all lysine and arginine residues in blue. **(E)** Microtubule co-pelleting assay with Lis1^WT^ (light grey) and Lis1^5A^ (dark grey). n = 3 replicates per condition. Statistical analysis was performed using an extra sum-of-squares F test; p<0.0001.

The first possibility predicts that Lis1 monomers would have no effect on dynein’s velocity. To test this, we measured the velocity of dynein in the presence of either dimeric or monomeric Lis1. Both dimers and monomers of Lis1 could slow dynein motility in a dose-dependent manner (Figure 3C), in agreement with a previous study (Huang et al., 2012). Thus, if Lis1 crosslinks dynein to microtubules, it must do so in the context of a monomer, with one face of Lis1 interacting with dynein and a different one interacting with microtubules. From a structural perspective, this is quite unlikely given that the angle between dynein and the microtubule would have to be ~15°, a stalk angle that is rarely (~1% of the population) observed experimentally (Can et al., 2019).

A prediction of this second model is that mutants that disrupt Lis1 binding to dynein would not disrupt Lis1 microtubule binding as the interfaces involved must be different. To test this, we used a Lis1 mutant where five amino acids (four of which are basic amino acids) that face site_ring_ were mutated to alanine (Figure 3D); we refer to this mutant as “Lis1^5A^”. We previously showed that Lis1^5A^ does not bind to dynein (Toropova et al., 2014). Here, using a microtubule pelleting assay, we found that Lis1^5A^ does not bind to microtubules either (Figure 3E), suggesting that the interaction between Lis1 and microtubules uses the same face of Lis1’s β-propeller. Therefore, a Lis1 monomer would not be capable of crosslinking dynein to microtubules. This, along with the finding that human Lis1 does not interact with microtubules (Htet et al., 2020), suggests that the binding of yeast Lis1 to microtubules is unlikely to play a role in its regulation of dynein. Rather, we propose that this is a nonspecific interaction between some basic amino acids in yeast Lis1 and negatively charged microtubules.

### Identification of Lis1 mutations that specifically disrupt regulation at site_stalk_

After interrogating the interaction between Lis1 and site_ring_, we turned to site_stalk_. Previously, we identified amino acids in dynein’s stalk (E3012, Q3014, and N3018) that were required for the Lis1-mediated decrease in dynein’s affinity for microtubules (DeSantis et al., 2017). However, the resolution of our previous structure was not high enough to accurately dock in the Lis1 ß-propeller at this site. Our new structure allowed us to build models and revealed this interaction in molecular detail, showing a new contact between Lis1 and dynein at site_stalk_ involving a short loop in AAA4 of dynein (Figure 1F and Figure 4A). To determine if this contact plays a role in dynein regulation by Lis1, we mutated Ser248 in the endogenous copy of Lis1 to Gln to introduce a steric clash with the loop in dynein’s AAA4 (Figure 4A). In a nuclear segregation assay, cells expressing Lis1^S248Q^ at wild type levels (Figure 2–figure supplement 1) showed a defect in nuclear segregation (Figure 4B). Thus, this new contact at site_stalk_ between AAA4 of dynein and Lis1 is also required for Lis1’s ability to regulate dynein in vivo.

**Figure 4.**
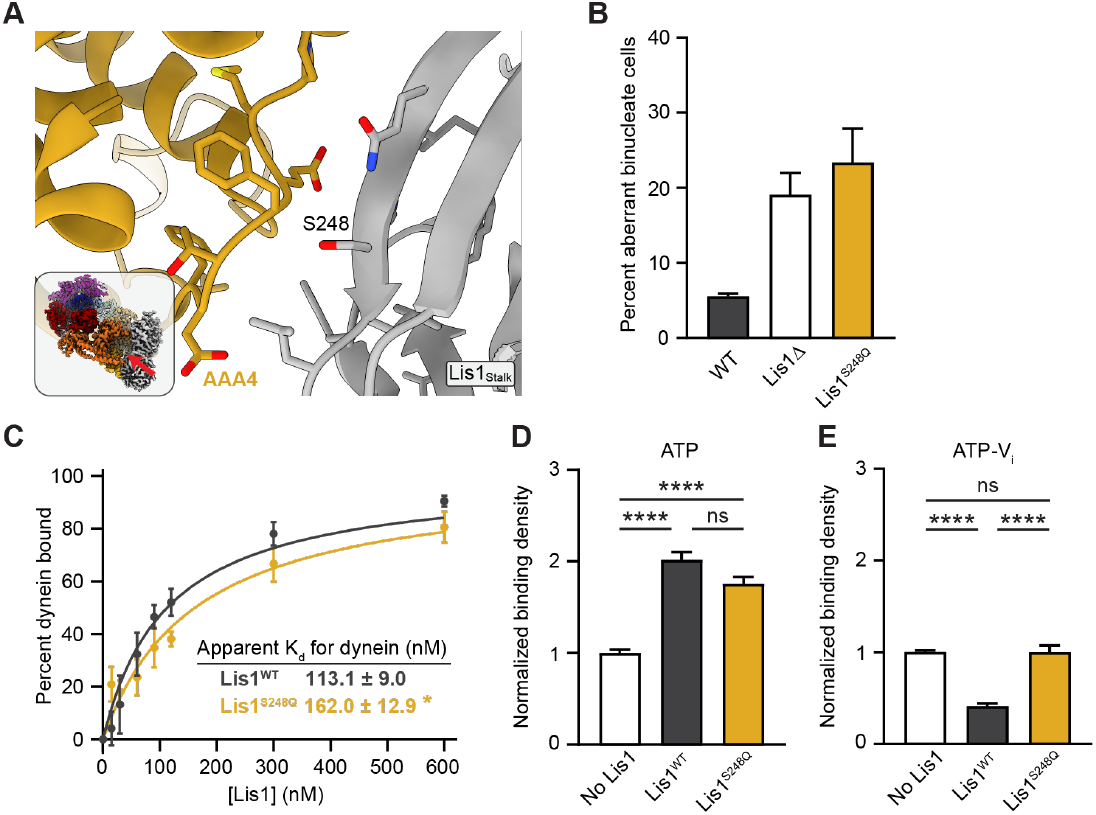
Lis1 regulation of dynein at site_stalk_. **(A)** Close-up view of the Lis1-dynein interface at site_stalk_. The residues at the interface are shown and the S248 mutated in this study is labeled. The inset shows the location of the close-up in the full structure. **(B)** Quantitation of the percentage of cells (mean ± s.e.m.) displaying an aberrant binucleate phenotype for WT (dark grey), Lis1 deletion (white) and Lis1^S248Q^ (yellow). n=3 replicates of at least 200 cells each per condition. **(C)** Quantitation of the apparent binding affinity of dynein for Lis1^WT^ (dark grey; K_d_ = 113.1 ± 9.0) and Lis1^S248Q^ (yellow; K_d_ = 162.0 ± 12.9). n = 3 replicates per condition. Statistical analysis was performed using an extra sum-of-squares F test; p = 0.0022. **(D)** Binding density (mean ± s.e.m.) of dynein on microtubules with ATP in the absence (white) or presence of Lis1^WT^ (dark grey) or Lis1^S248Q^ (yellow). Data was normalized to a density of 1.0 in the absence of Lis1. Statistical analysis was performed using an ANOVA; ****, p<0.0001; ns, p = 0.0614. n = 12 replicates per condition. **(E)** Binding density (mean ± s.e.m.) of dynein on microtubules with ATP-V_i_ in the absence (white) or presence of Lis1^WT^ (dark grey) or Lis1^S248Q^ (yellow). Data was normalized to a density of 1.0 in the absence of Lis1. Statistical analysis was performed using an ANOVA; ****, p<0.0001; ns, p > 0.9999. n = 12 replicates per condition.

We next probed the role of this new contact *in vitro*. We found that Lis1^S248Q^ showed a moderate decrease in binding affinity for dynein monomers (Figure 4C). Lis1^S248Q^ remained capable of inducing tight microtubule binding in the presence of ATP (regulation at site_ring_) (Figure 4D), but failed to decrease microtubule binding in the presence of ATP-V_i_ (loss of regulation at site_stalk_) (Figure 4E). Thus, Lis1 makes multiple contacts with dynein at site_stalk_ and Lis1^S248Q^ is the first Lis1 mutant capable of specifically targeting regulation at site_stalk_.

### The interaction between the two Lis1 ß-propellers is required for regulation at site_stalk_

The resolution of our new structure allowed us to build models for Lis1, revealing the interface between the two Lis1 ß-propellers when they are bound to dynein at both site_ring_ and site_stalk_ (Figure 5A). The Lis1–Lis1 interface consists mostly of hydrophobic interactions. To test if the interface is required for dynein regulation, we aimed to disrupt it by introducing charge clashes using F185D and I189D mutations in Lis1. Arg494 was also mutated to Ala to remove the only hydrogen bond donor/acceptor in the interface. In a nuclear segregation assay, Lis1^F185D, I189D, R494A^, which is expressed at wild type levels (Figure 2–figure supplement 1), resulted in a binucleate cell phenotype that was comparable to that seen with deletion of Lis1 (Figure 5B). Thus, the interaction between Lis1 β-propellers is required for Lis1’s ability to regulate dynein in vivo.

**Figure 5.**
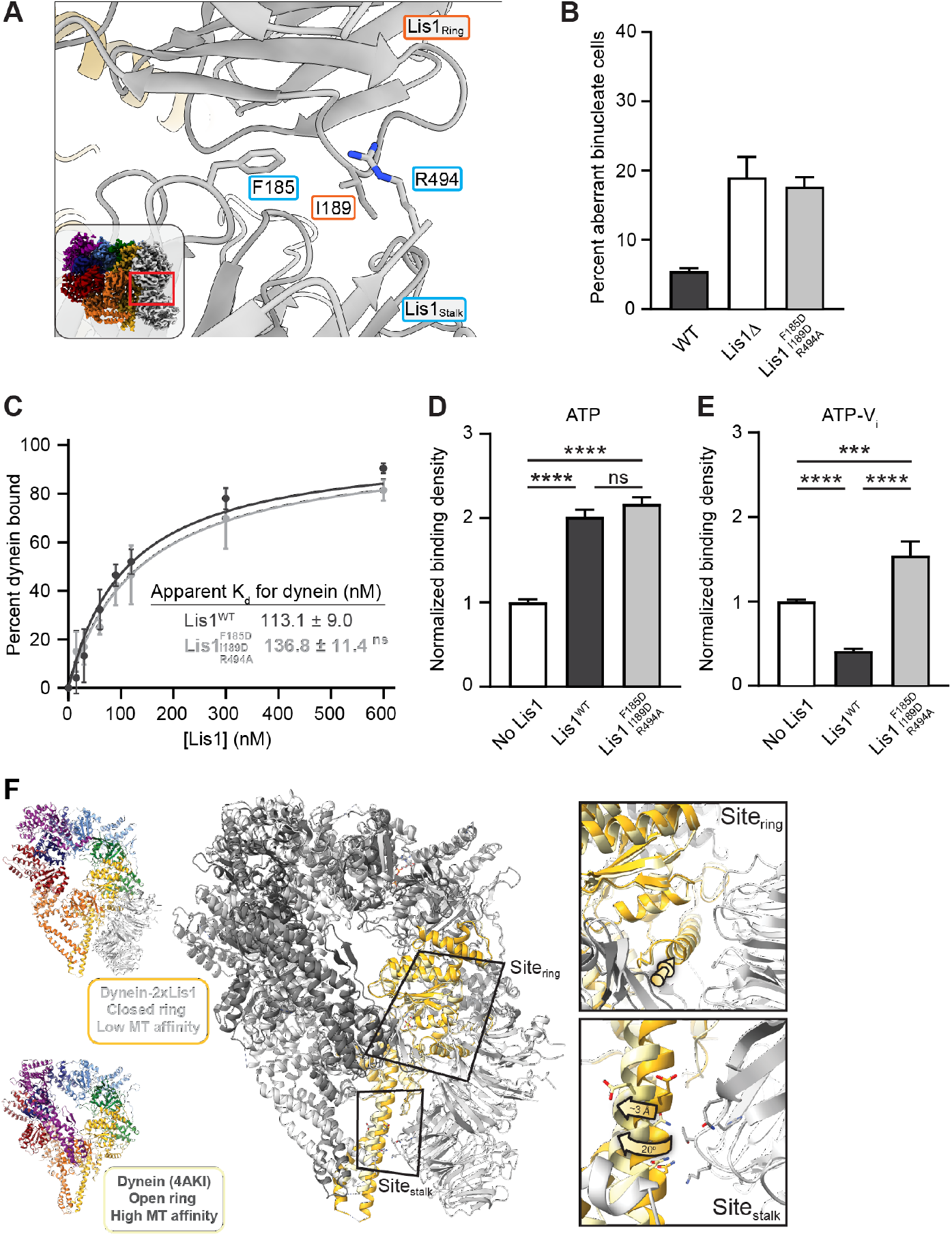
Role of the Lis1_ring_-Lis1_stalk_ interaction in Lis1’s regulation of dynein. **(A)** Close-up of the interface between the two Lis1 ß-propellers. The residues mutated in this study are indicated. The inset shows the location of the close-up in the full structure. **(B)** Quantitation (mean ± s.e.m.) of the percentage of cells displaying an aberrant binucleate phenotype for WT (dark grey), ΔLis1 (white) and Lis1^F185D, I189D, R494A^ (light grey). n=3 replicates of at least 200 cells each per condition. **(C)** Quantitation of the binding affinity of dynein for Lis1^WT^ (dark grey; K_d_ = 113.1 ± 9.0) and Lis1^F185D, I189D, R494A^ (light grey; K_d_ = 136.8 ± 11.4). n = 3 replicates per condition. Statistical analysis was performed using an extra sum-of-squares F test; p = 0.0974. Binding affinity of dynein for Lis1^WT^ repeated from figure 2. **(D)** Binding density (mean ± s.e.m.) of dynein on microtubules with ATP in the absence (white) or presence of Lis1^WT^ (dark grey) or Lis1^F185D, I189D, R494A^ (light grey). Data was normalized to a density of 1.0 in the absence of Lis1. Statistical analysis was performed using an ANOVA; ****, p<0.0001; ns, p = 0.4445. n = 12 replicates per condition. **(E)** Binding density (mean ± s.e.m.) of dynein on microtubules with ATP-V_i_ in the absence (dark grey) or presence of Lis1^WT^ (white) or Lis1^F185D, I189D, R494A^ (light grey). Data was normalized to a density of 1.0 in the absence of Lis1. Statistical analysis was performed using an ANOVA; ****, p<0.0001; ***, p = 0.0002. n = 12 replicates per condition. **(F)** Comparison between the structures of the dynein–(Lis1)_2_ complex (this work), where dynein’s ring is in its closed conformation (low affinity for the microtubule), and of dynein with its ring in the open conformation (high affinity for the microtubule) (PDB: 4AKI) (Schmidt et al., 2012) (left). The two structures were aligned using the globular portion of AAA4, where site_ring_ is located, and the two Lis1 ß-propellers from the dynein-(Lis1)_2_ structure are shown. Close-ups of site_ring_ and site_stalk_ (right) show that while the alpha helix to which Lis1 binds at site_ring_ is in a very similar position in both structures (top, yellow arrow), site_stalk_ has shifted by ~3Å and rotated by 20° away from where Lis1_Stalk_ would be located (bottom).

*In vitro*, Lis1^F185D, I189D, R494A^ has an affinity for dynein comparable to that of wild type Lis1 (Figure 5C). In microtubule binding assays this mutant showed no defects in its ability to stimulate tight microtubule binding by dynein in the presence of ATP (regulation at site_ring_) (Figure 5D). In contrast, not only did Lis1^F185D, I189D, R494A^ lose its ability to decrease dynein’s affinity for microtubules in the presence of ATP-V_i_ (loss of regulation at site_stalk_), it increased it (Figure 5E). This behavior was most similar to what we had previously observed when we mutated dynein’s site_stalk_ (E3012A, Q3014A, N3018A), a mutation that also led to a Lis1-dependent increase in microtubule binding in the presence of ATP-V_i_ (DeSantis et al., 2017). Thus, our data show that regulation of dynein at site_stalk_ requires both binding at site_stalk_ and the interaction between Lis1’s ß-propellers.

Given the importance of the Lis1–Lis1 interaction for Lis1’s regulation of dynein at site_stalk_, we wondered what prevents Lis1 from binding there when dynein’s ring is in its open conformation (high affinity for the microtubule), as our data showed that this interaction plays no role in Lis1’s regulation at site_ring_. A comparison of our structure of dynein–(Lis1)_2_ (with dynein’s ring in its closed conformation) with that of dynein in an open-ring conformation (PDB: 4AKI) (Schmidt et al., 2012) showed that while site_ring_ is very similar in both structures, site_stalk_ has shifted ~3Å and rotated 20° away from where Lis1_stalk_ would be located (Figure 5F). This suggests that the interaction between the two Lis1 ß-propellers prevents Lis1_stalk_ from reaching its binding site when dynein is in its open-ring state (high affinity for the microtubule).

### Dynein–Lis1 and Lis1–Lis1 contacts are required for dynein to reach the cortex

Next, we wanted to determine how Lis1 binding to dynein at site_ring_ and site_stalk_, as well as the Lis1–Lis1 interface contributed to dynein’s localization in yeast cells (Figure 6A and B). Yeast dynein is active at the cell cortex, where it pulls on spindle pole body (SPB)-anchored microtubules to position the mitotic spindle (Adames and Cooper, 2000). Dynein reaches the cortex by localizing to microtubule plus ends, either via kinesin-dependent transport or recruitment from the cytosol (Carvalho et al., 2004; Caudron et al., 2008; Markus et al., 2009). Lis1 is required for dynein’s localization to SPBs, microtubule plus ends, and the cell cortex (Lee et al., 2003; Li et al., 2005; Markus et al., 2009; Markus and Lee, 2011; Sheeman et al., 2003). To interrogate dynein’s localization, the endogenous copy of dynein was labeled at its carboxy terminus with 3xGFP in cells with fluorescently labeled microtubules (*TUB1-CFP*) and SPBs (*SPC110-tdTomato*) (Markus et al., 2009) (Figure 6C). We then determined the number of dynein foci per cell that were found at SPBs (Figure 6D), microtubule plus ends (Figure 6E), or the cell cortex (Figure 6F). As observed previously, deletion of Lis1 causes a significant decrease in dynein foci at all three sites (Figure 6D–F). Mutation of site_ring_ (Lis1^E253A, H254A^) also showed a significant decrease in dynein foci at all three sites and was not significantly different than the Lis1 deletion strain (Figure 6D–F). Mutation of site_stalk_ (Lis1^S248Q^) or the Lis1-Lis1 interface (Lis1^F185D, I189D, R494A^) showed intermediate phenotypes for dynein localization to SPBs or the cortex (Figure 6D and F).

**Figure 6.**
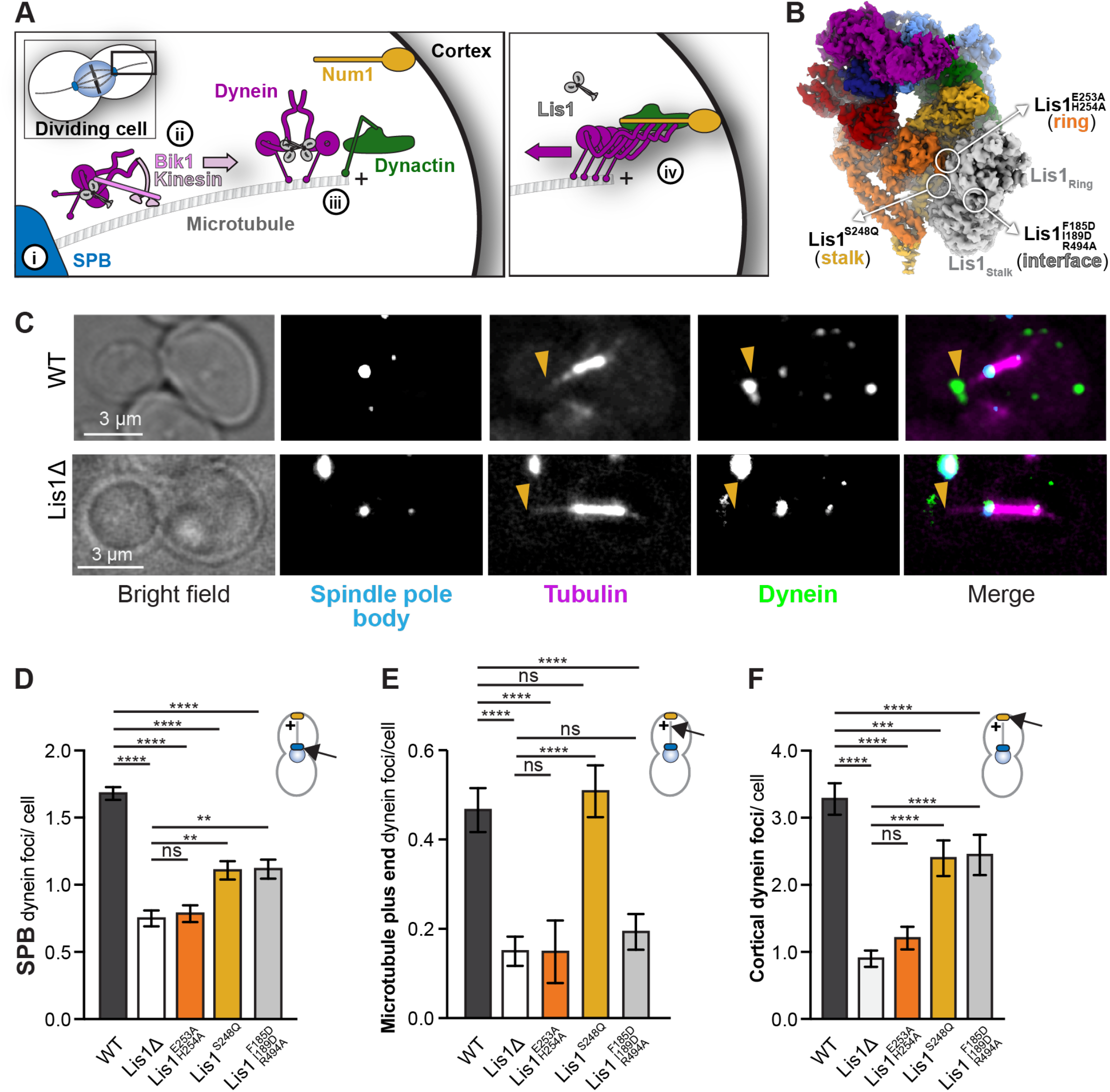
Lis1 binding sites on dynein have different roles in the localization of active dynein complexes. **(A)** Schematic showing the steps involved in targeting dynein to the cell cortex in a dividing yeast cell. Dynein is localized to the spindle pole body (SPB) (i), transported to the microtubule plus end by a kinesin (Kip2) in a Lis1-dependent manner (ii), maintained at the microtubule plus end in a Lis1-dependent manner (iii), and off-loaded to the cell cortex, which is also dependent on Lis1 (iv). Cortex-associated dynein is active and pulls on SPB-attached microtubules to position the mitotic spindle. **(B)** Our structure showing the location of the mutations that disrupt Lis1 binding at site_ring_, site_stalk_ and the Lis1:Lis1 interface. **(C)** Example images of dynein localization in dividing yeast cells. Orange arrowheads point to microtubule plus ends. **(D, E, F)** Quantification of dynein localization. Bar graphs show the average number of dynein foci per cell localized to the SPB (D), microtubule plus end (E) and cortex (F) in wild type, Lis1Δ, Lis1^E253A, H254A^, Lis1^S248Q^ and Lis1^F185D, I189D, R494A^ yeast strains. Statistical analysis was performed using a Kruskal-Wallis test; ****, p<0.0001; ***, p = 0.0001; ns, p > 0.9999. n = 120 cells per condition.

From a mechanistic perspective an important finding from this analysis came from comparing each mutant’s effects on dynein localization to the microtubule plus end versus its localization to the cortex (Figure 6E and F). Binding of Lis1 at site_ring_, but not site_stalk_, was required for microtubule plus end localization, suggesting that plus end localization requires the tight microtubule binding induced by Lis1 at site_ring_ (Figure 2F). In contrast, all three contacts are required for reaching the cortex, the site where fully assembled dynein is active to walk on microtubules. Overall, our analysis of these new Lis1 mutants supports a role for site_ring_, site_stalk_ and the Lis1–Lis1 interface in dynein complex assembly, which is initiated when dynein is bound tightly to microtubule plus ends.

### Site_ring_ and site_stalk_ are important for Lis1’s role in assembling active human dynein complexes

Our new structure allowed us to fully dissect the consequences of Lis1 binding to dynein at site_ring_ and site_stalk_ on dynein’s mechanochemistry and in vivo function in yeast. Next, we wondered whether these same sites were important for dynein complex formation. This analysis is most easily done using human proteins, where Lis1 enhances the formation of dynein–dynactin–activating adaptor complexes that contain two dynein dimers, which move faster than complexes containing a single dynein dimer (Elshenawy et al., 2020; Grotjahn et al., 2018; Htet et al., 2020; Urnavicius et al., 2018) (Figure 1A). To determine if Lis1 binding to dynein at site_ring_ and site_stalk_ was important for forming these fast-moving complexes, we mutated human dynein at both site_ring_ (K2898A, E2902G, E2903S and E2904G) and site_stalk_ (E3196A, Q3198A and N3202A); we will refer to this mutant as dynein^mut^. We purified wild type and mutant human dynein from baculovirus-infected insect cells expressing all of the dynein chains (Schlager et al., 2014) and measured the affinity of dynein^WT^ and dynein^mut^ for human Lis1. For these experiments, purified Halo-tagged Lis1 was covalently coupled to magnetic beads, mixed with purified human dynein, and the percentage of dynein bound to the beads was quantified. Lis1 bound to dynein^mut^ with an apparent K_d_ that was nearly three-fold weaker than that for dynein^WT^ (Figure 7B). The fact that these mutations do not completely abolish the dynein–Lis1 interaction raises that possibility that there are additional sites of dynein–Lis1 interaction and/ or that there are additional contacts between dynein and Lis1 and site_ring_ and site_stalk_ not seen in yeast.

**Figure 7.**
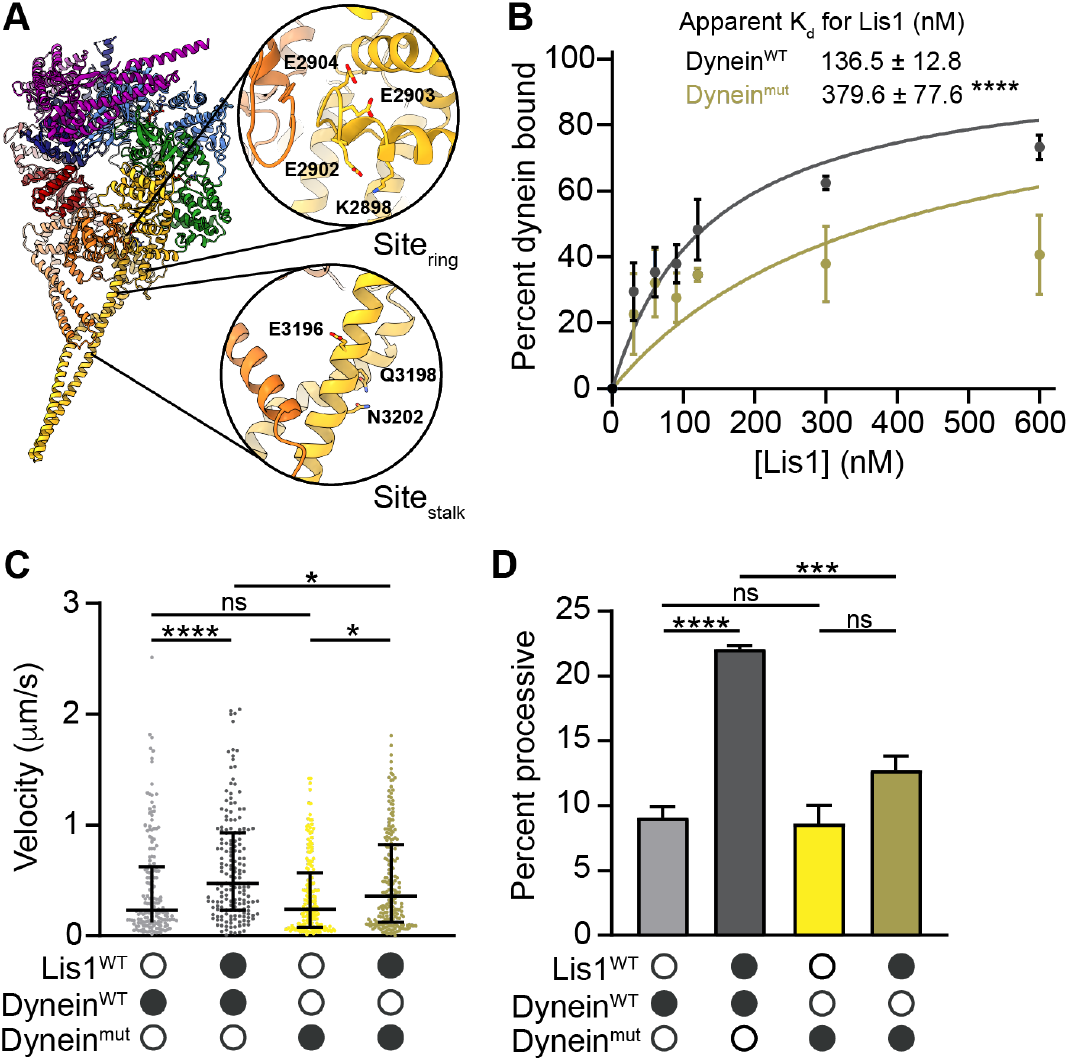
Site_ring_ and site_stalk_ are important for Lis1’s ability to increase dynein velocity. **(A)** Structure of the human dynein motor domain (PDB 5NUG) (Zhang et al., 2017) highlighting residues mutated at site_ring_ and site_stalk_. **(B)** Quantitation of the binding affinity of human Lis1 for dynein^WT^ (dark grey; K_d_ = 136.5 ± 12.8) and Dynein^mut^ (dark yellow; K_d_ = 379.6 ± 77.6). n = 3 replicates per condition. Statistical analysis was performed using an extra sum-of-squares F test; p = 0.0001. **(C)** Single-molecule velocity of human dynein–dynactin–Hook3 (DDH) complexes formed with dynein^WT^ (shades of grey) or dynein^mut^ (shades of yellow) in the absence or presence of 300 nM Lis1. Black circles indicate presence of a component in the assay; while white circles indicate the absence of a component. The median and interquartile range are shown. Statistical analysis was performed with a Kruskal-Wallis test; ****, p<0.0001; *, dynein^mut^ in the presence and absence of Lis1 p = 0.0190; *, dynein^WT^ and dynein^mut^ in the presence of Lis1, p = 0.0231; ns, p > 0.9999. At least 150 singlemolecule events were measured per condition. **(D)** Percentage (mean ± s.e.m.) of processive runs of dynein–dynactin–Hook3 complexes formed with dynein^WT^ (shades of grey) or dynein^mut^ (shades of yellow) in the absence or presence of 300 nM Lis1. n = 3 replicates per condition with each replicate including at least 200 individual single-molecule events. Black circles indicate presence of a component in the assay; while white circles indicate the absence of a component. Statistical analysis was performed with an ANOVA; ****, p < 0.0001; ***, p = 0.0008; ns, dynein^mut^ in the presence and absence of Lis1, p = 0.0755; ns, dynein^WT^ and dynein^mut^ in the absence of Lis1, p = 0.9883.

We next examined the single-molecule motility properties of dynein^WT^ and dynein^mut^, complexed with human dynactin and the activating adaptor Hook3. Complexes containing dynein^mut^ moved at the same velocity and showed the same percentage of processive runs when compared to dynein^WT^ in the absence of Lis1 (Figure 7C and D). As was shown previously (Elshenawy et al., 2020; Htet et al., 2020), the velocity of dynein^WT^ was significantly increased by Lis1 (Figure 7C), as were the number of processive runs (Figure 7D). In contrast, dynein^mut^’s velocity was only modestly increased by Lis1 (Figure 7C) and the percentage of processive runs did not significantly increase (Figure 7D). We observed a similar trend with dynein–dynactin complexes activated by BicD2 (Figure 7–figure supplement 1A). These results mirror our previous findings using a Lis1 mutant containing five point mutations (Lis1^5A^: R212A, R238A, R316A, W340A, K360A) (Htet et al., 2020). Here, we repeated this experiment using the same proteins we used for the dynein^mut^ experiments and find that Lis1^5A^ and dynein^mut^ have similar defects in Lis1 regulation of dynein (Figure 7–figure supplement 1B and C). These data indicate that site_ring_ and site_stalk_ are important for Lis1’s role in forming the activated human dynein–dynactin–activating adaptor complex.

## Discussion

Previous structures of yeast dynein–Lis1 complexes (DeSantis et al., 2017; Toropova et al., 2014) were not of high enough resolution to reliably fit homology models based on the human Lis1 structure (Tarricone et al., 2004). Here we report the first high-resolution structure of cytoplasmic dynein-1 bound to Lis1, which allowed us to build an atomic model (Figure 1). The structure revealed the molecular details of the interactions between the Lis1 ß-propellers and its two binding sites on dynein as well as how the two ß-propellers interact with each other. Our previous work had identified the main binding sites on dynein (DeSantis et al., 2017), but not their molecular details or their counterparts on Lis1. All of these can be clearly seen in the current structure. The much higher resolution of the structure also revealed additional contacts between Lis1 and dynein that were not apparent in previous cryo-EM maps; these involve interactions between Lis1_ring_ and a loop in AAA5 (Figure 1F) and Lis1_stalk_ and a loop in AAA4 (Figure 1G). Finally, being able to visualize (and mutate) the interface between the two Lis1 ß-propellers allowed us to test its role in the regulation of dynein by Lis1. Critical to achieving high resolution was the use of specialized streptavidin affinity grids (Han et al., 2016, 2012; Lahiri et al., 2019), which help overcome preferred orientation of the sample, a major resolution-limiting factor in cryo-EM.

We used our new structure to guide mutagenesis of both yeast and human proteins to probe the role of Lis1 in regulating dynein’s mechanochemistry and role in assembling dynein–dynactin–activating adaptor complexes. Mutations at all of these contact sites in yeast showed that all are important for the mechanochemistry of *S. cerevisiae* dynein. Mutations in both dynein and Lis1 showed that multiple contact sites at site_ring_ are important for Lis1’s ability to induce dynein to tightly bind microtubules, which reduces dynein’s velocity. On the other hand, mutations of Lis1 at site_stalk_ and the Lis1–Lis1 interface showed that those interactions are important for Lis1’s other main effect on dynein’s mechanochemistry: the weakening of dynein’s interactions with microtubules when AAA3 is bound to ATP (ADP-V_i_ was used here experimentally).

Mutations at all of these sites in yeast also revealed that they are important for dynein’s in vivo function in mitotic spindle positioning, which ultimately leads to nuclear segregation following mitosis. Further analysis of these mutants demonstrated that binding of Lis1 at site_ring_, but not site_stalk_, was required for microtubule plus end localization. Given that Lis1 binding at site_ring_ increases dynein’s affinity for microtubules, this suggests that Lis1-induced tight microtubule binding is required for dynein’s localization to microtubule plus ends (Figure 8A, panel ii). This is in agreement with a study suggesting that dynein’s motor domain, but not its tail domain, is required for localization to microtubule plus ends (Markus et al., 2009). We also found that Lis1 binding to dynein at both site_ring_ and site_stalk_, as well as the Lis1–Lis1 interface was required for dynein to reach the cell cortex, the site where active yeast dynein–dynactin complexes interact with the candidate activating adaptor, Num1 (Figure 8A, panel iii) (Heil-Chapdelaine et al., 2000; Lammers and Markus, 2015; Lee et al., 2003; Sheeman et al., 2003).

**Figure 8.**
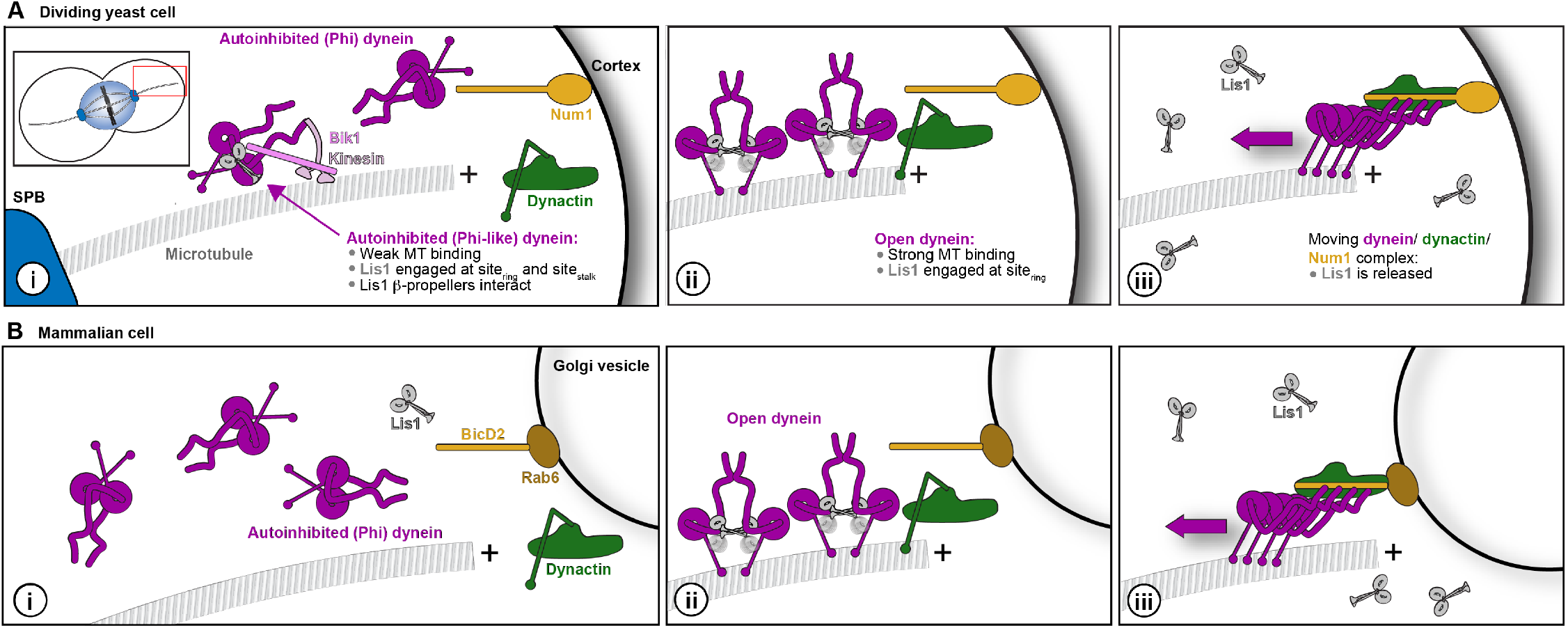
Model for a conserved role of Lis1 in dynein regulation in yeast and human cells. **(A and B)** Model of dynein regulation by Lis1 in *S. cerevisiae* (A) and mammalian cells (B). In *S. cerevisiae*, dynein arrives at microtubule plus ends either via transport by kinesin in a complex that contains Lis1 or by recruitment from the cytoplasm (i). Dynein that is recruited from the cytoplasm is most likely in a Phi conformation. Once at microtubule plus ends, Lis1 binding would favor an open conformation. Our mutant analysis suggests that Lis1 induces tight microtubule binding here (ii). Finally, dynein interacts with the candidate activating adaptor, Num on the cortex and Lis1 is released. We hypothesize that a similar pathway operates in mammalian cells. In our model we show the activating adaptor BicD2 binding to Rab6 on Golgi-derived vesicles (Hoogenraad and Akhmanova, 2016).

Our structure of a dynein–Lis1 complex shows that binding of Lis1 to dynein at site_ring_ is sterically incompatible with formation of the autoinhibited Phi conformation of dynein (Htet et al., 2020; Zhang et al., 2017). In addition, yeast dynein mutants that are defective in forming the Phi conformation partially bypass the requirement for Lis1 in reaching the cortex (Marzo et al., 2020). Thus, we propose that Lis1 binding to dynein at site_ring_, site_stalk_ and the Lis1–Lis1 interface shifts the equilibrium from the Phi to a (partially) open conformation of dynein (Figure 8A, panels i and iii), ultimately allowing dynein complexes to fully assemble with dynactin and Num1 (Figure 8A, panel iii).

A similar pathway for dynein regulation may exist in other organisms, including mammalian cells (Figure 8B), and related mechanisms have recently been proposed (Markus et al., 2020; Qiu et al., 2021). Our work shows that both the site_ring_ and site_stalk_ Lis1 binding sites on dynein are important for the formation of active human dynein–dynactin–activating adaptor complexes containing either the Hook3 or BicD2 activating adaptors. Previously, we and others found that Lis1 was also important for the formation of activated dynein complexes containing the activating adaptors Ninl and BicDL1 (also called BicDR1) (Elshenawy et al., 2020; Htet et al., 2020), thus it is likely that this role for Lis1 and both binding sites on dynein will be required for dynein complex formation driven by other activating adaptors. Overall, this work and that of others (Elshenawy et al., 2020; Htet et al., 2020; Markus et al., 2020; Marzo et al., 2020; Qiu et al., 2019, n.d.) suggests that the field is coalescing around a unified model for dynein regulation of Lis1 across species.

## Acknowledgments

We thank the Nikon Imaging Center at UC San Diego for advice on imaging and analysis and the Cryo-EM Facility at UC San Diego. We also thank our funding sources: JMR is a Merck Fellow of the Damon Runyon Cancer Research Foundation, DRG-2370-19; JPG was funded by the Molecular Biophysics Training Grant, NIH Grant T32 GM008326; EPK is funded by a Jane Coffin Childs Postdoctoral Fellowship; AL’s lab by NIH R01 GM107214; and SRP’s lab by the Howard Hughes Medical Institute and NIH R35 GM141825. The authors have no competing financial interests.

## Materials and Methods

### Yeast Strain Construction

The *S. cerevisiae* strains used in this study are listed in Table S1. The endogenous genomic copies of *DYN1* and *PAC1* (encoding the dynein heavy chain and Lis1, respectively) were modified or deleted using PCR-based methods as previously described (Longtine et al., 1998). Transformations were performed using the lithium acetate method and screened by colony PCR. Point mutants were generated using QuikChange site-directed mutagenesis (Agilent) and verified by DNA sequencing.

### Nuclear segregation assay

Log-phase S. cerevisiae cells growing at 30°C were transferred to 16°C for 16 hr. Cells were fixed with 75% ethanol and stained with DAPI. Imaging was performed using a 100x Apo TIRF NA 1.49 objective on a Nikon Ti2 microscope with a Yokogawa-X1 spinning disk confocal system, MLC400B laser engine (Agilent), Prime 95B back-thinned sCMOS camera (Teledyne Photometrics), and a piezo Z-stage (Mad City Labs). The percentage of aberrant binucleate cells was calculated as the number of binucleate cells divided by the sum of wild-type and binucleate cells.

### Cloning, plasmid construction, and mutagenesis

The pDyn1 plasmid (the pACEBac1 expression vector containing insect cell codon-optimized dynein heavy chain (*DYNC1H1*) fused to a His-ZZ-TEV tag on the amino-terminus and a carboxy-terminal SNAPf tag (New England Biolabs)) and the pDyn2 plasmid (the pIDC expression vector with codon optimized *DYNC1I2*, *DYNC1LI2*, *DYNLT1*, *DYNLL1*, and *DYNLRB1*) were recombined in vitro with a Cre recombinase (New England Biolabs) to generate the pDyn3 plasmid. The presence of all six dynein chains was verified by PCR. pDyn1, pDyn2 and the pFastBac plasmid with codon-optimized human full-length Lis1 (*PAFAH1B1*) fused to an amino-terminal His-ZZ-TEV tag was a gift from Andrew Carter (LMB-MRC, Cambridge, UK). The Hook3 construct contained amino acids 1-552 and was obtained as described previously (Redwine et al., 2017). Activating adaptors were fused to a ZZ-TEV-HaloTag (Promega) on the amino-terminus and inserted into a pET28a expression vector. All additional tags were added via Gibson assembly and all mutations and truncations were made via site-directed mutagenesis (Agilent).

### S. cerevisiae protein purification

Protein purification steps were done at 4°C unless otherwise indicated. Dynein constructs were purified from *S. cerevisiae* using a ZZ tag as previously described (Reck-Peterson et al., 2006). Briefly, liquid nitrogen-frozen yeast cell pellets were lysed by grinding in a chilled coffee grinder and resuspended in dynein lysis buffer (DLB: 30 mM HEPES [pH 7.4], 50 mM potassium acetate, 2 mM magnesium acetate, 1 mM EGTA, 10% glycerol, 1 mM DTT) supplemented with 0.1 mM Mg-ATP, 0.5 mM Pefabloc, 0.05% Triton and cOmplete EDTA-free protease inhibitor cocktail tablet (Roche). The lysate was clarified by centrifuging at 264,900 x g for 1 hr or at 125,100 x g for 2 hr. The clarified supernatant was incubated with IgG Sepharose beads (GE Healthcare Life Sciences) for 1 hr. The beads were transferred to a gravity flow column, washed with DLB buffer supplemented with 250 mM potassium chloride, 0.1 mM Mg-ATP, 0.5 mM Pefabloc and 0.1% Triton, and with TEV buffer (10 mM Tris–HCl [pH 8.0], 150 mM potassium chloride, 10% glycerol, 1 mM DTT, and 0.1 mM Mg-ATP). GST-dimerized dynein constructs were labeled with 5 mM Halo-TMR (Promega) in the column for 10 min at room temperature and the unbound dye was washed away with TEV buffer at 4 °C. Dynein was cleaved from IgG beads via incubation with 0.15 mg/mL TEV protease (purified in the Reck-Peterson lab) for 1 hr at 16°C. For dynein monomer constructs, the TEV cleavage step was done overnight at 4°C and the cleaved proteins were concentrated using 100K MWCO concentrator (EMD Millipore). Cleaved proteins were filtered by centrifuging with Ultrafree-MC VV filter (EMD Millipore) in a tabletop centrifuge and flash frozen in liquid nitrogen.

Lis1 constructs were purified from S. cerevisiae using their His_8_ and ZZ tags as previously described (Huang et al., 2012). Lysis and clarification steps were similar to dynein purification except for the lysis buffer used was buffer A (50 mM potassium phosphate [pH 8.0], 150 mM potassium acetate, 150 mM sodium chloride, 2mM magnesium acetate, 5mM b-mercaptoethanol, 10% glycerol, 0.2% Triton, 0.5 mM Pefabloc) supplemented with 10 mM imidazole (pH 8.0) and cOmplete EDTA-free protease inhibitor cocktail tablet. The clarified supernatant was incubated with Ni-NTA agarose (QIAGEN) for 1 hr. The Ni beads were transferred to a gravity column, washed with buffer A + 20 mM imidazole (pH 8.0) and eluted with buffer A + 250 mM imidazole (pH 8.0). The eluted protein was incubated with IgG Sepharose beads for 1 hr. IgG beads were transferred to a gravity flow column, washed with buffer A + 20 mM imidazole (pH 8.0) and with modified TEV buffer (50 mM Tris–HCl [pH 8.0], 150mM potassium acetate, 2 mM magnesium acetate, 1 mM EGTA, 10% glycerol, 1 mM DTT). Lis1 was cleaved from the IgG beads via incubation with 0.15 mg/mL TEV protease (purified in the Reck-Peterson lab) for 1 hr at 16°C. Cleaved proteins were filtered by centrifuging with Ultrafree-MC VV filter (EMD Millipore) in a tabletop centrifuge and flash frozen in liquid nitrogen.

### Human protein purification

Human full-length dynein, human dynein monomer, and human Lis1 constructs were expressed in Sf9 cells as described previously (Baumbach et al., 2017; Htet et al., 2020; Schlager et al., 2014). Briefly, the pDyn3 plasmid containing the human dynein genes or the pFastBac plasmid containing full-length Lis1 was transformed into DH10EmBacY chemically competent cells with heat shock at 42°C for 15 seconds followed by incubation at 37°C for 5 hours in S.O.C media (Thermofisher scientific). The cells were then plated on LB-agar plates containing kanamycin (50 μg/ml), gentamicin (7 μg/ml), tetracycline (10 μg/ml), BluoGal (100 μg/ml) and IPTG (40 μg/ml) and positive clones were identified by a blue/white color screen after 48 hours. For full-length human dynein constructs, white colonies were additionally tested for the presence of all six dynein genes using PCR. These colonies were then grown overnight in LB medium containing kanamycin (50 μg/ml), gentamicin (7 μg/ml) and tetracycline (10 μg/ml) at 37°C. Bacmid DNA was extracted from overnight cultures using an isopropanol precipitation method as described previously (Zhang et al., 2017). 2mL of Sf9 cells at 0.5×10^6^ cells/mL were transfected with 2μg of fresh bacmid DNA and FuGene HD transfection reagent (Promega) at a 3:1 transfection reagent to DNA ratio according to the manufacturer’s instructions. After three days, the supernatant containing the “V0” virus was harvested by centrifugation at 200 x g for 5 minutes at 4°C. To generate “V1”, 1 mL of the V0 virus was used to transfect 50mL of Sf9 cells at 1×10^6^ cells/mL. After three days, the supernatant containing the V1 virus was harvested by centrifugation at 200 x g for 5 minutes at 4°C and stored in the dark at 4°C until use. For protein expression, 4 mL of the V1 virus were used to transfect 400 mL of Sf9 cells at 1×10^6^ cells/mL. After three days, the cells were harvested by centrifugation at 3000 x g for 10 minutes at 4°C. The pellet was resuspended in 10 mL of ice-cold PBS and pelleted again. The pellet was flash frozen in liquid nitrogen and stored at −80°C.

Protein purification steps were done at 4°C unless otherwise indicated. Full-length dynein and dynein monomer were purified from frozen Sf9 pellets transfected with the V1 virus as described previously (Schlager et al., 2014). Frozen cell pellets from a 400 mL culture were resuspended in 40 mL of Dynein-lysis buffer (50 mM HEPES [pH 7.4], 100 mM sodium chloride, 1 mM DTT, 0.1 mM Mg-ATP, 0.5 mM Pefabloc, 10% (v/v) glycerol) supplemented with 1 cOmplete EDTA-free protease inhibitor cocktail tablet (Roche) per 50 mL and lysed using a Dounce homogenizer (10 strokes with a loose plunger and 15 strokes with a tight plunger). The lysate was clarified by centrifuging at 183,960 x g for 88 min in Type 70 Ti rotor (Beckman). The clarified supernatant was incubated with 4 mL of IgG Sepharose 6 Fast Flow beads (GE Healthcare Life Sciences) for 3-4 hours on a roller. The beads were transferred to a gravity flow column, washed with 200 mL of Dynein-lysis buffer and 300 mL of TEV buffer (50 mM Tris–HCl [pH 8.0], 250 mM potassium acetate, 2 mM magnesium acetate, 1 mM EGTA, 1 mM DTT, 0.1 mM Mg-ATP, 10% (v/v) glycerol). For fluorescent labeling of carboxy-terminal SNAPf tag, dynein-coated beads were labeled with 5 μM SNAP-Cell-TMR (New England Biolabs) in the column for 10 min at room temperature and unbound dye was removed with a 300 mL wash with TEV buffer at 4°C. The beads were then resuspended and incubated in 15 mL of TEV buffer supplemented with 0.5 mM Pefabloc and 0.2 mg/mL TEV protease (purified in the Reck-Peterson lab) overnight on a roller. The supernatant containing cleaved proteins was concentrated using a 100K MWCO concentrator (EMD Millipore) to 500 μL and purified via size exclusion chromatography on a TSKgel G4000SWXL column (TOSOH Bioscience) with GF150 buffer (25 mM HEPES [pH7.4], 150 mM potassium chloride, 1mM magnesium choloride, 5 mM DTT, 0.1 mM Mg-ATP) at 1 mL/min. The peak fractions were collected, buffer exchanged into a GF150 buffer supplemented with 10% glycerol, concentrated to 0.1-0.5 mg/mL using a 100K MWCO concentrator (EMD Millipore) and flash frozen in liquid nitrogen.

Lis1 constructs were purified from frozen cell pellets from 400 mL culture. Lysis and clarification steps were similar to full-length dynein purification except Lis1-lysis buffer (30 mM HEPES [pH 7.4], 50 mM potassium acetate, 2 mM magnesium acetate, 1 mM EGTA, 300 mM potassium chloride, 1 mM DTT, 0.5 mM Pefabloc, 10% (v/v) glycerol) supplemented with 1 cOmplete EDTA-free protease inhibitor cocktail tablet (Roche) per 50 mL was used. The clarified supernatant was incubated with 0.5 mL of IgG Sepharose 6 Fast Flow beads (GE Healthcare Life Sciences) for 2-3 hours on a roller. The beads were transferred to a gravity flow column, washed with 20 mL of Lis1-lysis buffer, 100 mL of modified TEV buffer (10 mM Tris–HCl [pH 8.0], 2 mM magnesium acetate, 150mM potassium acetate, 1 mM EGTA, 1 mM DTT, 10% (v/v) glycerol) supplemented with 100 mM potassium acetate, and 50 mL of modified TEV buffer. For fluorescent labeling of Lis1 constructs with amino-terminal HaloTags, Lis1-coated beads were labeled with 200 μM Halo-TMR (Promega) for 2.5 hours at 4°C on a roller and the unbound dye was removed with a 200 mL wash with modified TEV buffer supplemented with 250 mM potassium acetate. Lis1 was cleaved from IgG beads via incubation with 0.2 mg/mL TEV protease overnight on a roller. The cleaved Lis1 was filtered by centrifuging with an Ultrafree-MC VV filter (EMD Millipore) in a tabletop centrifuge and flash frozen in liquid nitrogen.

Dynactin was purified from stable HEK293T cell lines expressing p62-Halo-3xFlag as described previously (Redwine et al., 2017). Briefly, frozen pellets collected from 160 x 15 cm plates were resuspended in 80 mL of Dynactin-lysis buffer (30 mM HEPES [pH 7.4], 50 mM potassium acetate, 2 mM magnesium acetate, 1 mM EGTA, 1 mM DTT, 10% (v/v) glycerol) supplemented with 0.5 mM Mg-ATP, 0.2% Triton X-100 and 1 cOmplete EDTA-free protease inhibitor cocktail tablet (Roche) per 50 mL and rotated slowly for 15 min. The lysate was clarified by centrifuging at 66,000 x g for 30 min in Type 70 Ti rotor (Beckman). The clarified supernatant was incubated with 1.5 mL of anti-Flag M2 affinity gel (Sigma-Aldrich) overnight on a roller. The beads were transferred to a gravity flow column, washed with 50 mL of wash buffer (Dynactin-lysis buffer supplemented with 0.1 mM Mg-ATP, 0.5 mM Pefabloc and 0.02% Triton X-100), 100 mL of wash buffer supplemented with 250 mM potassium acetate, and again with 100 mL of wash buffer. For fluorescent labeling the HaloTag, dynactin-coated beads were labeled with 5 μM Halo-JF646 (Janelia) in the column for 10 min at room temperature and the unbound dye was washed with 100 mL of wash buffer at 4°C. Dynactin was eluted from beads with 1 mL of elution buffer (wash buffer with 2 mg/mL of 3xFlag peptide). The eluate was collected, filtered by centrifuging with Ultrafree-MC VV filter (EMD Millipore) in a tabletop centrifuge and diluted to 2 mL in Buffer A (50 mM Tris-HCl [pH 8.0], 2 mM magnesium acetate, 1 mM EGTA, and 1 mM DTT) and injected onto a MonoQ 5/50 GL column (GE Healthcare and Life Sciences) at 1 mL/min. The column was pre-washed with 10 CV of Buffer A, 10 CV of Buffer B (50 mM Tris-HCl [pH 8.0], 2 mM magnesium acetate, 1 mM EGTA, 1 mM DTT, 1 M potassium acetate) and again with 10 CV of Buffer A at 1 mL/min. To elute, a linear gradient was run over 26 CV from 35-100% Buffer B. Pure dynactin complex eluted from ~75-80% Buffer B. Peak fractions containing pure dynactin complex were pooled, buffer exchanged into a GF150 buffer supplemented with 10% glycerol, concentrated to 0.02-0.1 mg/mL using a 100K MWCO concentrator (EMD Millipore) and flash frozen in liquid nitrogen.

Activating adaptors containing amino-terminal HaloTags were expressed in BL-21[DE3] cells (New England Biolabs) at OD 0.4-0.6 with 0.1 mM IPTG for 16 hr at 18°C. Frozen cell pellets from 2 L culture were resuspended in 60 mL of activator-lysis buffer (30 mM HEPES [pH 7.4], 50 mM potassium acetate, 2 mM magnesium acetate, 1 mM EGTA, 1 mM DTT, 0.5 mM Pefabloc, 10% (v/v) glycerol) supplemented with 1 cOmplete EDTA-free protease inhibitor cocktail tablet (Roche) per 50 mL and 1 mg/mL lysozyme. The resuspension was incubated on ice for 30 min and lysed by sonication. The lysate was clarified by centrifuging at 66,000 x g for 30 min in Type 70 Ti rotor (Beckman). The clarified supernatant was incubated with 2 mL of IgG Sepharose 6 Fast Flow beads (GE Healthcare Life Sciences) for 2 hr on a roller. The beads were transferred to a gravity flow column, washed with 100 mL of activator-lysis buffer supplemented with 150 mM potassium acetate and 50mL of cleavage buffer (50 mM Tris–HCl [pH 8.0], 150 mM potassium acetate, 2 mM magnesium acetate, 1 mM EGTA, 1 mM DTT, 0.5 mM Pefabloc, 10% (v/v) glycerol). The beads were then resuspended and incubated in 15 mL of cleavage buffer supplemented with 0.2 mg/mL TEV protease overnight on a roller. The supernatant containing cleaved proteins were concentrated using a 50K MWCO concentrator (EMD Millipore) to 1 mL, filtered by centrifuging with Ultrafree-MC VV filter (EMD Millipore) in a tabletop centrifuge, diluted to 2 mL in Buffer A (30 mM HEPES [pH 7.4], 50 mM potassium acetate, 2 mM magnesium acetate, 1 mM EGTA, 10% (v/v) glycerol and 1 mM DTT) and injected onto a MonoQ 5/50 GL column (GE Healthcare and Life Sciences) at 1 mL/min. The column was pre-washed with 10 CV of Buffer A, 10 CV of Buffer B (30 mM HEPES [pH 7.4], 1 M potassium acetate, 2 mM magnesium acetate, 1 mM EGTA, 10% (v/v) glycerol and 1 mM DTT) and again with 10 CV of Buffer A at 1 mL/min. To elute, a linear gradient was run over 26 CV from 0-100% Buffer B. The peak fractions containing Halo-tagged activating adaptors were collected and concentrated to using a 50K MWCO concentrator (EMD Millipore) to 0.2 mL. For fluorescent labeling the HaloTag, the concentrated peak fractions were incubated with 5 μM Halo-Alexa488 (Promega) for 10 min at room temperature. Unbound dye was removed by PD-10 desalting column (GE Healthcare and Life Sciences) according to the manufacturer’s instructions. The labeled activating adaptor sample was concentrated using a 50K MWCO concentrator (EMD Millipore) to 0.2 mL, diluted to 0.5 mL in GF150 buffer and further purified via size exclusion chromatography on a Superose 6 Increase 10/300 GL column (GE Healthcare and Life Sciences) with GF150 buffer at 0.5 mL/min. The peak fractions were collected, buffer exchanged into a GF150 buffer supplemented with 10% glycerol, concentrated to 0.2-1 mg/mL using a 50K MWCO concentrator (EMD Millipore) and flash frozen in liquid nitrogen.

### Electron microscopy sample preparation

Dynein was buffer exchanged into Modification Buffer (100 mM sodium phosphate, pH 8.0, 150 mM sodium chloride) and biotinylated using water soluble Sulfo ChromaLink biotin (Sulfo ChromaLINK Biotin, Cat #B-1007) in a 1:2 molar ratio for 2 hr at room temperature. Biotinylated protein was dialyzed into TEV buffer overnight at 4 °C. Biotinylation of dynein was verified using a pull-down assay with streptavidin magnetic beads (Thermo Fisher).

Streptavidin affinity grids (SA grids) were prepared as previously described (Han et al., 2016). Just prior to use, the SA grids were washed by touching the SA side of the grid to three drops (2x 50 μL, 1 x 100 μL) of rehydration buffer (150 mM potassium chloride, 50 mM HEPES pH 7.4, 5 mM EDTA) to remove the storage trehalose layer. For complete removal of the protective trehalose layer, rehydration buffer was manually pipetted up and down onto the SA side of the grid, and then the grid was placed onto a 100 μL drop of rehydration buffer for 10 minutes. This process was repeated once, and then the grid was buffered exchanged into sample buffer using five 50 μL drops of sample buffer (TEV buffer without glycerol and DTT, supplemented with 1.2 mM ATP and 1.2 mM Na_3_VO_4_). 4 μL of sample (150 nM dynein, 650 nM Lis1, 1.2 mM ATP, 1.2 mM Na_3_VO_4_) was applied to each grid and incubated for ~10 minutes inside a humidity chamber. Unbound protein was washed away by touching the sample side of the grid to three 100 μL drops of wash buffer supplemented with 150 nM Lis1 (26 mM Tris pH 8, 75 mM potassium chloride, 64 mM potassium acetate, 0.85 mM magnesium acetate, 0.5 mM EGTA, 1.2 mM ATP, 1.2 mM Na_3_VO_4_). Grids were vitrified using a Vitrobot (FEI) set to 20°C and 100% humidity. Vitrobot tweezers were kept cold on ice between samples. The sample was first manually wicked inside the Vitrobot using a Whatman No. 1 filter paper, followed by the addition of 1.5 μL of sample buffer before standard blotting and vitrification proceeded (Blot time: 4s and blot force: 20).

### Image collection and processing

Vitrified grids were imaged on a Titan Krios (FEI) operated at an accelerating voltage of 300 kV and the images were recorded with a K2 Summit direct electron detector (Gatan Inc.). We collected 2378 movies in super resolution mode (0.655 Å/super-resolution pixel at the object level) dose-fractionated into 200 ms frames for a total exposure of 10 s with a dose rate of 10 electrons/pixel/s for a total fluence of 58.3 electrons/Å^2^. The defocus of the images varied within a range of −2.0 μm to −2.7 μm. Automated data collection was executed by SerialEM (Mastronarde, 2005).

Movie frames were aligned using the dose-weighted frame alignment option in UCSF MotionCor2 (Zheng et al., 2017) as employed in Relion 3.0 (Zivanov et al., 2018). At this stage the individual frames were corrected for the anisotropic magnification distortion inherent to the electron microscope (Grant and Grigorieff, 2015). The signal from the streptavidin lattice was removed from aligned micrographs using Fourier filtering as described previously (Han et al., 2012). Micrographs were manually inspected for defects including sub-optimal ice thickness and incomplete removal of the SA lattice signal and 46 micrographs were removed leaving 2332 micrographs for further processing. CTF estimation was carried out on the non-dose weighted micrographs using GCTF (Zhang, 2016) using the local CTF estimation option as implemented in Appion (Lander et al., 2009). Images with CTF fits having 0.5 confidence resolution worse than 5 Å were excluded from further processing. Particles were picked using crYOLO (Wagner et al., 2019) using a training model generated from manually picked particles. The particles were extracted with a downsampled pixel size of 3.93 Å/pixel, and a single round of two-dimensional (2D) classification were carried out in Relion 3.0 to identify bad particles. Particles belonging to good 2D class averages were recentered and extracted (1.31 Å/pixel), and further processed in cryoSPARC. Ab initio models were generated with the majority of particles going into one good class displaying the characteristic features of the dynein motor domain and the WD40 rings of Lis1. Those particles were refined using the non-uniform refinement routine of cryoSPARC (Punjani et al., 2020), against the best ab initio model. This resulted in a 3.2 Å map containing 233 476 particles. We performed per-particle CTF refinement and beam tilt refinement in Relion 3.0 followed by a single round of three-dimensional (3D) classification without alignment. The particles contributing to one of the classes lead to a high resolution reconstruction of the dynein-Lis1 complex and those particles were further refined in Relion 3.0 to generate the final 3.1 Å map (based on the 0.143 cutoff of the gold-standard FSC curve) of the dynein-Lis1 complex. For visualisation purposes the map was sharpened with an automatically estimated negative B-factor of 33 (as determined from the “PostProcess” routine of Relion 3.0). The local resolution of the map was estimated using the “local-resolution” routine of Relion 3.0 and the map was low pass filtered according to the local resolution prior to analysis. The 3-D FSC was calculated by 3dfsc version 2.5 (Tan et al., 2017) using the half-maps and the mask used for the PostProcess routine in Relion 3.0.

### Model building

The dynein-Lis1 map was segmented using Seggar (Pintilie et al., 2010) as implemented in UCSF chimera (Pettersen et al., 2004) and the atomic models of the different components were initially built to account for their respective segmented maps. A homology of the Lis1 WD40 domain was generated using I-TASSER (Roy et al., 2010) and this model was rigid body docked into the corresponding segmented map using the “fit in map” routing in UCSF Chimera. Regions of the model that did not agree with the EM map were manually rebuilt using Coot (Emsley et al., 2010). The resulting model was used as a reference for Rosetta CM (Song et al., 2013) and 1200 models were generated. A hybrid model was made using the two lowest energy output models from Rosetta CM. This model was then manually placed using Coot into both Lis1 sites, and areas of disagreement between the map and the model were resolved manually.

A homology model of the yeast dynein motor domain was created in Swiss-Model (Waterhouse et al., 2018) by using the human dynein atomic model in the closed conformation (PDB 5NUG) (Zhang et al., 2017) as a template. The resulting model was fit in the segmented map corresponding to the dynein motor domain using the “fit to map” command in UCSF Chimera, and manually rebuilt in Coot to improve the agreement between the atomic model and the map. The resulting model was used as a template for Rosetta CM and 700 models were generated. The lowest energy model was selected and manually placed into the map, followed by an additional round of manual rebuilding

The resulting models of the dynein motor domain and Lis1 WD40 domains were combined, and further refined against the unsegmented dynein-Lis1 map using an using an iterative process between Phenix Real Space Refine (Afonine et al., 2018) and manual rebuilding in Coot.

3D variable analysis (Punjani et al., 2020) was carried out in cryoSPARC using the particles and mask used in the non-uniform refinement job resulting in the 3.2 Å map (Figure 1-supplemental figure 1). Default parameters were used to solve 3 modes and results were filtered to 5 Å. Results were visualized using the 3D variability display using the simple output mode with 20 frames/cluster, and then viewed using ChimeraX (Pettersen et al., 2021).

### Binding curves

To assess dynein/Lis1 binding, Lis1 was first covalently coupled to 16 μL of SNAP-Capture Magnetic Beads (New England Biolabs) in 2 mL Protein Lo Bind Tubes (Eppendorf) using the following protocol. Beads were washed twice with 1 mL of modified TEV buffer. Lis1 (0-600 nM) was added to the beads and gently shaken for one hour. The supernatant was analyzed via SDS-PAGE to confirm complete depletion of Lis1. The Lis1-conjugated beads were washed once with 1 mL modified TEV buffer and once with 1mL of binding buffer (10 mM Tris–HCl [pH 8.0], 150 mM potassium chloride, 2 mM magnesium chloride, 10% glycerol, 1 mM DTT, 0.1% NP40, 1 mM ADP). 20 nM dynein (monomeric motor domain) diluted in binding buffer was added to the Lis1-conjugated beads and gently agitated for 30 minutes. The supernatant was analyzed via SDS-PAGE, stained with SYPRO^™^ Red (Thermo Fisher) and dynein depletion was determined using densitometry in ImageJ. Binding curves were fit in Prism8 (Graphpad) with a nonlinear regression for one site binding with Bmax set to 1.

### Single-molecule TIRF microscopy

Single-molecule imaging was performed with an inverted microscope (Nikon, Ti-E Eclipse) equipped with a 100x 1.49 N.A. oil immersion objective (Nikon, Plano Apo), a ProScan linear motor stage controller (Prior) and a LU-NV laser launch (Nikon), with 405 nm, 488 nm, 532 nm, 561 nm and 640 nm laser lines. The excitation and emission paths were filtered using appropriate single bandpass filter cubes (Chroma). The emitted signals were detected with an electron multiplying CCD camera (Andor Technology, iXon Ultra 897). Illumination and image acquisition was controlled by NIS Elements Advanced Research software (Nikon).

Single-molecule motility and microtubule binding assays were performed in flow chambers assembled as described previously (Case 1997) using the TIRF microscopy set up described above. Either biotin-PEG-functionalized coverslips (Microsurfaces) or No. 1-1/2 coverslips (Corning) sonicated in 100% ethanol for 10 min were used for the flow-chamber assembly. Taxol-stabilized microtubules with ~10% biotin-tubulin and ~10% fluorescent-tubulin (Alexa405- or 488-labeled) were prepared as described previously (Roberts et al., 2014). Flow chambers were assembled with taxol-stabilized microtubules by incubating sequentially with the following solutions, interspersed with two washes with assay buffer (30 mM HEPES [pH 7.4], 2 mM magnesium acetate, 1 mM EGTA, 10% glycerol, 1 mM DTT) supplemented with 20 μM Taxol and 50 mM potassium acetate in between: (1) 1 mg/mL biotin-BSA in assay buffer (3 min incubation, ethanol washed coverslips only); (2) 1 mg/mL streptavidin in assay buffer (3 min incubation) and (3) a fresh dilution of taxol-stabilized microtubules in assay buffer (3 min incubation). After flowing in microtubules, the flow chamber was washed twice with assay buffer supplemented with 1 mg/mL casein and 20 μM Taxol.

For assays using *S. cerevisiae* proteins dynein was incubated with Lis1 or modified TEV buffer (to buffer match for Lis1) in assay buffer supplemented with 50 mM potassium acetate for 10 minutes before flowing into the assembled flow chamber. The final assay buffer was supplemented with 1 mg/mL casein, 71.5 mM β-mercaptoethanol, an oxygen scavenger system (0.4% glucose, 45 mg/ml glucose catalase, and 1.15 mg/ml glucose oxidase), and 2.5 mM Mg-ATP. For experiments in the presence of vanadate 2.5 mM sodium vanadate was included. The final concentration of dynein was 2-15 pM. For single-molecule microtubule binding assays, the final imaging mixture containing dynein was incubated for an additional 5 min in the flow chamber at room temperature before imaging. After 5 min incubation, microtubules were imaged first by taking a single-frame snapshot. Dynein was imaged by taking a single-frame snapshot. Each sample was imaged at 4 different fields of view and there were between 5 and 10 microtubules in each field of view. In order to compare the effect of Lis1 on microtubule binding, the samples with and without Lis1 were imaged in two separate flow chambers made on the same coverslip on the same day with the same stock of polymerized tubulin as described previously (Roberts et al., 2014). For singlemolecule motility assays, microtubules were imaged first by taking a single-frame snapshot. Dynein was imaged every 1s for 5 min. At the end, microtubules were imaged again by taking a snapshot to assess stage drift. Movies showing significant drift were not analyzed. Each sample was imaged no longer than 15 min.

To assemble human dynein-dynactin-activating adaptor complexes, purified dynein (10-20 nM concentration), dynactin and the activating adaptor were mixed at 1:2:10 molar ratio and incubated on ice for 10 min. These dynein-dynactin-activating adaptor complexes were then incubated with Lis1 or modified TEV buffer (to buffer match for experiments without Lis1) for 10 min on ice. The mixtures of dynein, dynactin, activating adaptor or dynein alone and Lis1 were then flowed into the flow chamber assembled with taxol-stabilized microtubules. The final imaging buffer contained the assay buffer with 20 μM Taxol, 1 mg/mL casein, 7.5 mM potassium chloride, 71.5 mM β-mercaptoethanol, an oxygen scavenger system, and 2.5 mM Mg-ATP. The final imaging buffer also contained 60mM potassium acetate for experiments using Hook3 as the activating adaptor and 30mM potassium acetate for experiments with BicD2. The final concentration of dynein in the flow chamber was 0.5-1. The final concentration of Lis1 was between 12 nM - 300 nM (as indicated in the main text). Imaging was performed as above except with images taken every 300 msec for 3 min.

### Single-molecule microtubule binding assay analysis

Intensity profiles of dynein spots from a single-frame snapshot were generated over a 5-pixel wide line drawn perpendicular to the long axis of microtubules in ImageJ. Intensity peaks at least 2-fold higher than the neighboring background intensity were counted as dynein bound to microtubules. Bright aggregates that were 5-fold brighter than the neighboring intensity peaks were not counted. The average binding density was calculated as the total number of dynein spots divided by the total microtubule length in each snapshot. Normalized binding density was calculated by dividing by the average binding density of dynein without Lis1 collected on the same coverslip (see above). Data plotting and statistical analyses were performed in Prism8 (GraphPad).

### Single-molecule motility assay analysis

Kymographs were generated from motility movies and dynein velocity was calculated from kymographs using ImageJ macros as described (Roberts et al., 2014). Only runs that were longer than 4 frames (4 s) were included in the analysis. Bright aggregates, which were less than 5% of the population, were excluded from the analysis. Data plotting and statistical analyses were performed in Prism8 (GraphPad).

### Microtubule co-pelleting assay

Unlabeled Taxol-stabilized MTs were polymerized as above and free tubulin was removed by centrifugation through a 60% glycerol gradient in BRB80 (80 mM PIPES-KOH pH 6.8, 1 mM magnesium chloride, 1 mM EGTA, 1 mM DTT, 20 μM Taxol) for 15min at 100000xg and 37°C. The MT pellet was resuspended in DLB supplemented with 20 μM Taxol. MTs (0-600nM tubulin) were incubated with 100 nM Lis1 for 10 minutes before being pelleted for 15 min at 100000xg and 25°C. The supernatant was analyzed via SDS-PAGE and depletion was determined using densitometry in ImageJ. Binding curves were fit in Prism8 (GraphPad) with a nonlinear regression for one site binding with Bmax set to 1.

## Supplemental Figures and Data

**Figure 1–figure supplement 1.**
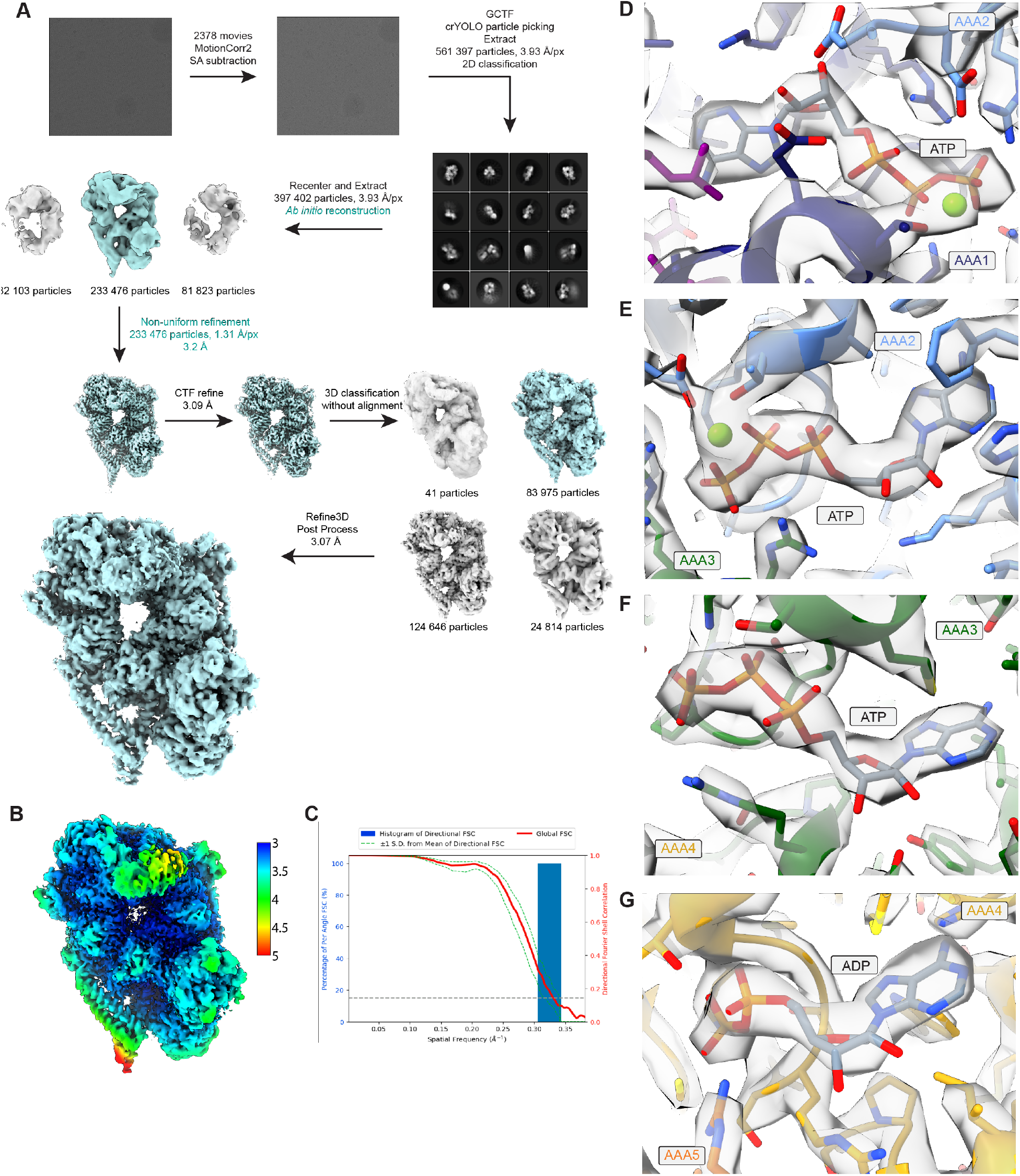
Details of cryo-EM data processing and nucleotide binding. **(A)** Workflow for cryo-EM data processing. Steps highlighted in teal were implemented in cryoSPARC while all others were carried out in Relion 3.0. **(B)** Cryo-EM map of the dynein-(Lis1)_2_ complex colored by local resolution. **(C)** Fourier shell correlation (FSC) plot. **(D-G)** Regions of the cryo-EM map corresponding to ATP in AAA1 (D), AAA2 (E), AAA3 (F), and ADP in AAA4 (G).

**Figure 1–figure supplement 2.**
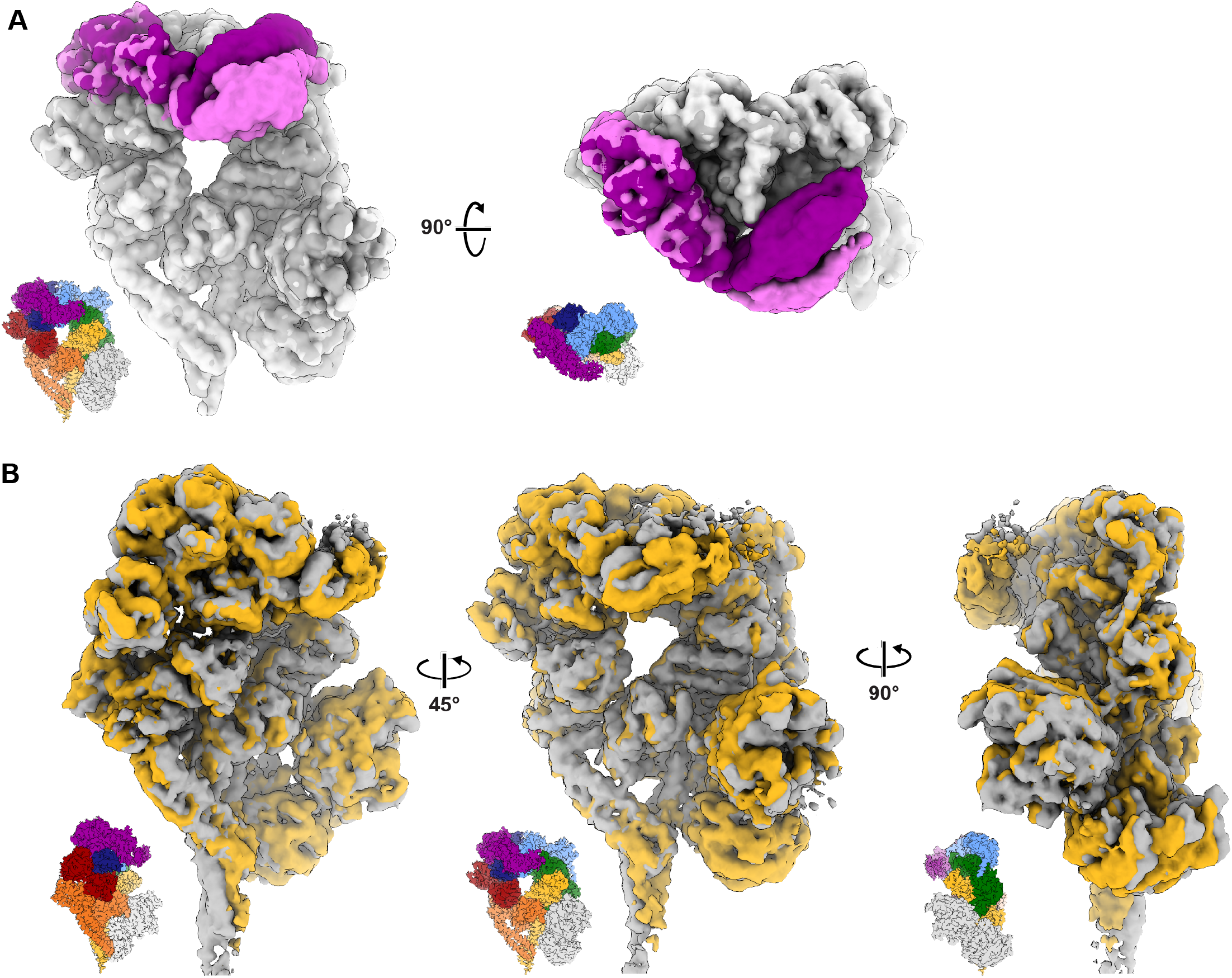
3D variability analysis of dynein–(Lis1)_2_. Volumes at the start and end of variability components 1 and 2 were calculated using the 3DVA display job in cryoSPARC. **(A)** The first variability component describes motion observed in the linker domain. The first and last volumes were colored light gray/ light purple and dark gray/ dark purple, respectively. **(B)** The second variability component reveals a subtle open and closing motion in the AAA ring with associated movements in the buttress and stalk helices. See Movie S2 as well.

**Figure 2–figure supplement 1.**
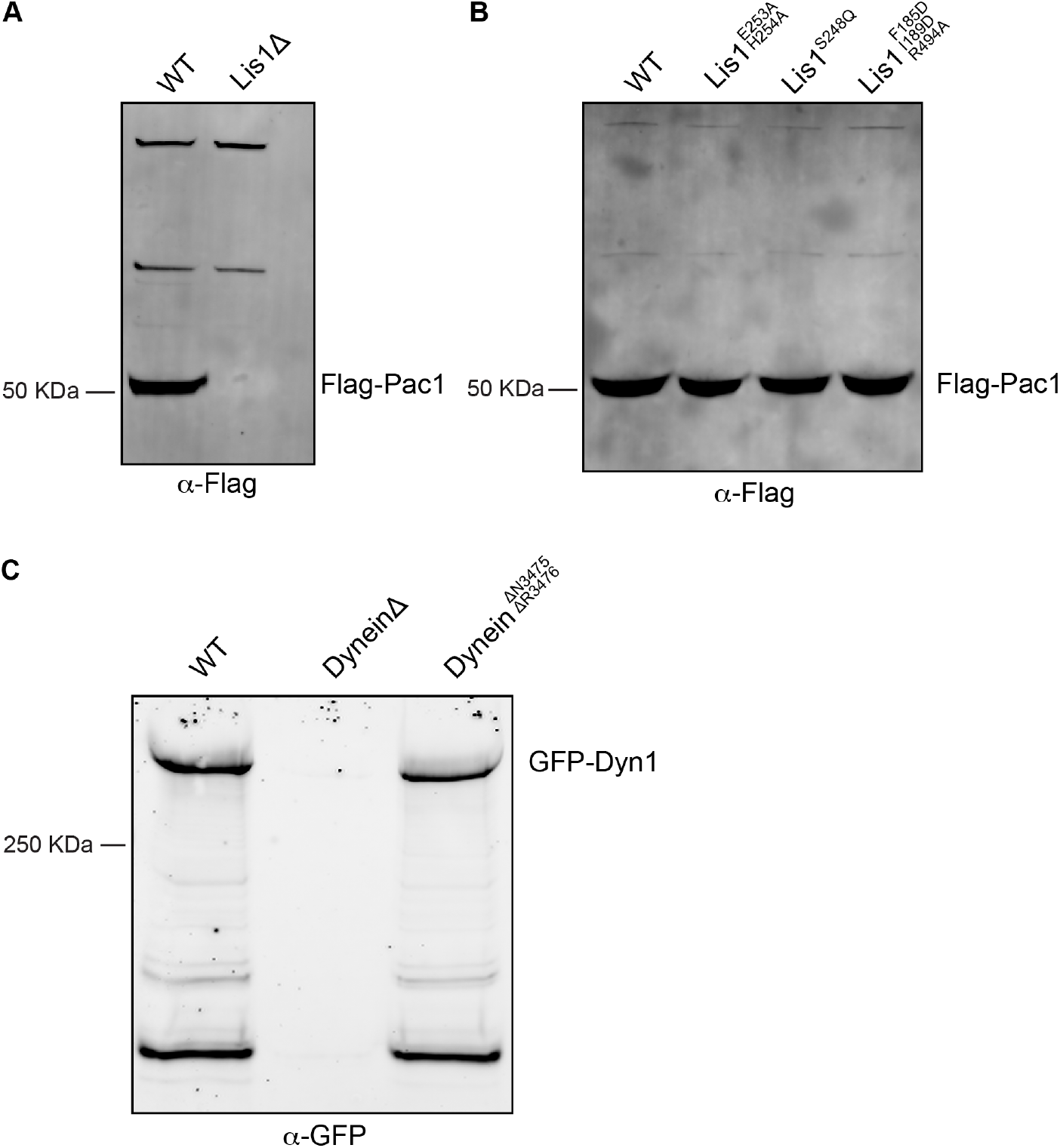
Western blot analysis of Lis1 and dynein mutant expression. **(A)** Cell lysates from wild type (WT) or Lis1Δ *S. cerevisiae* strains containing Flag-tagged Lis1 (Pac1) were immunoprecipitated using anti-Flag agarose beads and immunoblotted for the FLAG peptide. **(B)** Cell lysates from wild type (WT) or mutant Lis1 *S. cerevisiae* containing Flag-tagged Lis1 (Pac1) were immunoprecipitated using anti-FLAG agarose beads and immunoblotted for the FLAG peptide. **(C)** Cell lysates from wild type (WT), DyneinΔ and dynein^ΔN3475, ΔR3476^ *S. cerevisiae* strains containing GFP-dynein (Dyn1) were immunoprecipitated using anti-GFP nanobody beads and immunoblotted with an anti-GFP antibody.

**Figure 7–figure supplement 1.**
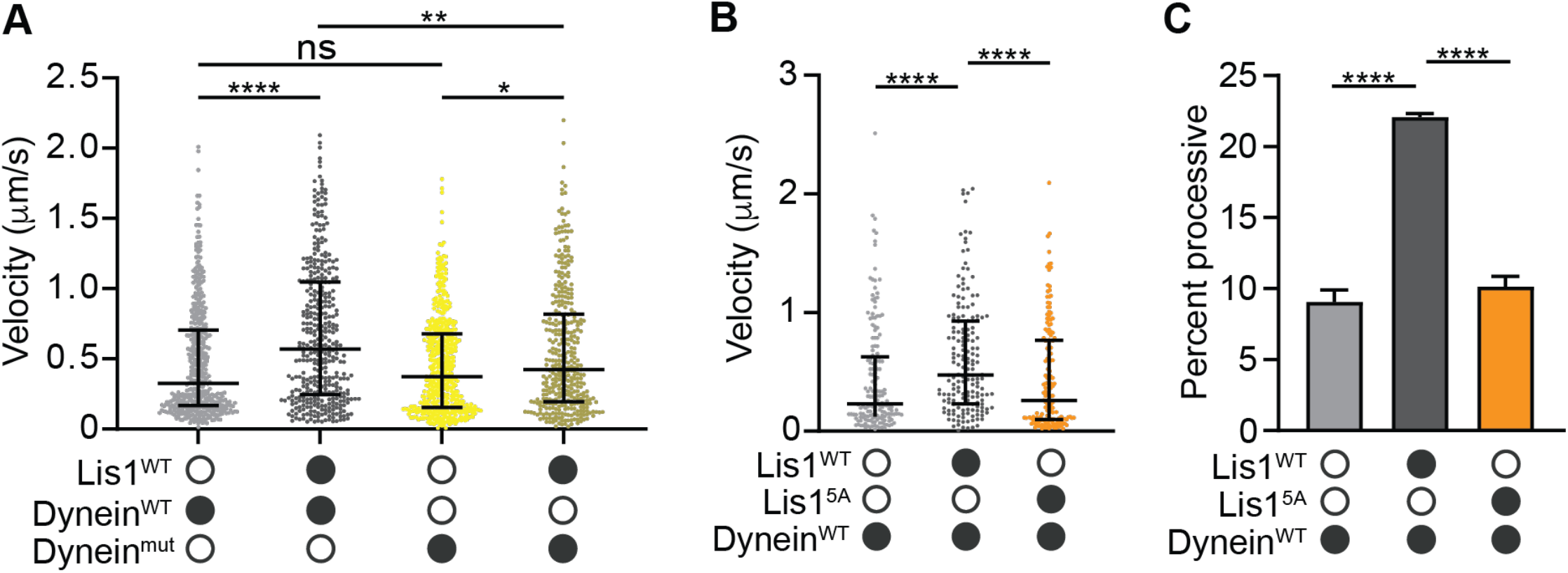
Mutations in Lis1 reduce Lis1’s ability to increase dynein’s velocity. **(A)** Single-molecule velocity of human dynein–dynactin–BicD2 complexes formed with dynein^WT^ (shades of grey) or dynein^mut^ (shades of yellow) in the absence or presence of 12.5 nM Lis1. The median and interquartile range are shown. Statistical analysis was performed with a Kruskal-Wallis test; ****, p<0.0001; **, p = 0.0055; *, p = 0.0175; ns, p > 0.9999. At least 350 single-molecule events were measured per condition. **(B)** Single-molecule velocity of dynein–dynactin–Hook3 complexes formed with dynein^WT^ in the absence or presence of 300 nM Lis1^WT^ or Lis1^5A^. The median and interquartile range are shown. Statistical analysis was performed with a Kruskal-Wallis test; ****, p<0.0001. At least 150 single molecule events were measured per condition. **(C)** Percentage (mean ± s.e.m.) of processive runs of dynein–dynactin–Hook3 complexes formed with dynein^WT^ in the absence or presence of 300 nM Lis1^WT^ or Lis1^5A^. n = 3 replicates per condition with each replicate including at least 200 individual single-molecule events. Statistical analysis was performed with an ANOVA; ****, p < 0.0001.

## Additional Supplementary Files

**Supplementary file 1. *S. cerevisiae* strains used in this work.** *DHA* and *SNAP* refer to the HaloTag (Promega) and SNAP-tag (NEB), respectively. TEV indicates a Tev protease cleavage site. *P_GAL1_* denotes the galactose promoter, which was used for inducing strong expression of Lis1 and dynein motor domain constructs. Amino acid spacers are indicated by g (glycine) and gs (glycine-serine).

**Supplementary file 2. Cryo-EM data collection parameters and model refinement statistics.**

## Movie Legends

**Movie 1. A 3.1Å structure of the dynein-Lis1 complex.**

Overview of the dynein–(Lis1)_2_ complex. The key interactions between dynein and Lis1 at site_ring_, and site_stalk_, as well as the Lis1-Lis1 interface, are highlighted.

**Movie 2. 3DVA analysis of Dynein bound to Lis1.**

3DVA analysis reveals conformational changes along the first two variability components. The first variability component describes motion attributed to the linker pulling away from AAA2. The second variability component describes changes in the overall motor domain as it undergoes subtle open and closing motions in the AAA ring.

