## Supplementary file 2 for "Structural Basis for Cytoplasmic Dynein-1 Regulation by Lis1"

### Cryo-EM data collection, refinement and validation statistics

| | Dynein AAA3 Walker B bound to Lis1 |
|---|---|
| **Models** | |
| PDB | 7MGM |
| EMDB | 23829 |
| **Data collection and processing** | |
| Microscope | Titan Krios |
| Camera | Gatan K2 Summit |
| Voltage (kV) | 300 |
| Electron exposure (e–/Å$^2$) | 58.3 |
| Magnification | |
| Defocus range (μm) | 2 – 2.7 |
| Pixel size (Å) | 1.31 |
| **Reconstruction** | |
| Symmetry imposed | C1 |
| Initial particle images (no.) | 561 397 |
| Final particle images (no.) | 83 975 |
| Micrographs collected (no.) | 2378 |
| Map resolution (Å) (0.143 FSC threshold) | 3.1 |
| **Model Refinement** | |
| Initial model used (PDB code) | 5NUG |
| Map-to-model resolution (0.5 FSC threshold) (Å) | 3.2 |
| Map sharpening *B* factor (Å$^2$) | 78.77 |
| *Model composition* | |
| Non-hydrogen atoms | 24 080 |
| Protein residues | 2964 |
| Ligands | 4 |
| *B* factors (Å$^2$) | |
| Protein | 74.23 |
| Ligand | 56.39 |
| *R.m.s. deviations* | |
| Bond lengths (Å) | 0.004 |
| Bond angles (°) | 0.481 |
| *Validation* | |
| MolProbity score | 1.40 |
| Clashscore | 5.74 |
| Poor rotamers (%) | 0.00 |
| *Ramachandran* | |
| Favored (%) | 97.60 |
| Allowed (%) | 2.40 |
| Disallowed (%) | 0.00 |
