## Supplementary file 1 for "Structural Basis for Cytoplasmic Dynein-1 Regulation by Lis1"

Supplementary file 1. *S. cerevisiae* strains used in this study.

| Strain | Genotype | Source |
|---|---|---|
| RPY1 | W303a (*MATa; his3-11,15; ura3-1; leu2-3,112; ade2-1; trp1-1*) | (Eshel et al., 1993) |
| RPY799 | W303a; *pep4Δ::HIS5; prb1Δ; dyn1Δ::CgLEU2; GAL1-8HIS-ZZ-SNAPgs-PAC1* | (DeSantis et al., 2017) |
| RPY816 | W303a; *pep4Δ::HIS5; prb1Δ; GAL1-8HIS-ZZ-Tev-PAC1; dyn1Δ::CgLEU2; ndl1Δ::HPH* | (DeSantis et al., 2017) |
| RPY1042 | W303a; *pep4Δ::HIS5; prb1Δ; GAL1-8HIS-ZZ-Tev-PAC1(aa 3-129 delete)-g-1XFLAG-gaSNAP-kanR; dyn1Δ::CgLEU2; ndl1Δ::HPH* | (Huang et al., 2012) |
| RPY1167 | W303a; *pep4Δ::HIS5; pGAL-ZZ-TEV-GFP-3XHA-GST-DYN1*$_{331kDa}$*-gsDHA-KanR; prb1Δ; pac1Δ; ndl1Δ::CgLEU2* | (DeSantis et al., 2017) |
| RPY1302 | W303a; *PGal:ZZ:Tev:DYN1*$_{331kDa}$ *pep4D::HIS5; prb1Δ PAC11-13Myc-TRP; pac1Δ::HPH* | (DeSantis et al., 2017) |
| RPY1547 | W303a; *pep4Δ::HIS5; prb1Δ; GAL1-8HIS-ZZ-Tev-PAC1(R275A,R301A,R378A,W419A,K437A); dyn1Δ::CgLEU2; ndl1Δ::HPH* | (Toropova et al., 2014) |
| RPY1717 | *MATa lys2-801; leu2-D1; his3-D200; trp1-D63; DYN1-3XGFP::TRP1; ura3-52::CFP-TUB1::URA3; SPC110-tdTomato::SpHIS5; ura3Δ::KanMX; pac1Δ::KlURA3* | (DeSantis et al., 2017) |
| RPY1749 | W303a; *pep4Δ::HIS5; prb1Δ; GAL1-8HIS-ZZ-Tev-PAC1(S248Q); dyn1Δ::CgLEU2; ndl1Δ::HPH* | This work |
| RPY1751 | W303a; *pep4Δ::HIS5; prb1Δ; GAL1-8HIS-ZZ-Tev-PAC1(E253A-H254A); dyn1Δ::CgLEU2; ndl1Δ::HPH* | This work |
| RPY1758 | W303a; *pep4Δ::HIS5; pGAL-ZZ-TEV-GFP-3XHA-GST-DYN1*$_{331kDa}$*(Δ AA3475-3476::URA3)-gsDHA-KanR; prb1Δ; pac1Δ; ndl1Δ::CgLEU2* | This work |
| RPY1760 | W303a; *pep4Δ::HIS5; prb1Δ; GAL1-8HIS-ZZ-Tev-PAC1(F185D-I189D-R494A); dyn1Δ::cgLEU2; ndl1Δ::HPH* | This work |
| RPY1790 | *W303a; PGal:ZZ:Tev:D6-Dyn1(Δ AA3475-3476); pep4Δ::HIS5; prb1Δ; PAC11-13Myc-TRP; pac1Δ::HPH* | This work |
| RPY1791 | W303a; *pep4Δ::HIS5; prb1Δ; GAL1-8HIS-ZZ-Tev-SNAPgs-PAC1(E253A-H254A); dyn1Δ::cgLEU2; ndl1Δ::HPH* | This work |
| RPY1792 | W303a; *pep4Δ::HIS5; prb1Δ; GAL1-8HIS-ZZ-Tev-SNAP-PAC1(F185D-I189D-R494A), dyn1Δ::cgLEU2; ndl1Δ::HPH* | This work |
| RPY1793 | W303a; *pep4Δ::HIS5; prb1Δ; GAL1-8HIS-ZZ-Tev-SNAP-PAC1(S248Q); dyn1Δ::CgLEU2; ndl1Δ::HPH* | This work |

| | | |
|---|---|---|
| RPY1795 | *MATa; lys2-801; leu2-D1; his3-D200; trp1-D63; DYN1-3GFP::TRP1; ura3-52::CFP-TUB1::URA3; SPC110-tdTomato::SpHIS5; Flag-PAC1; ura3Δ::KanMX* | This work |
| RPY1796 | *MATa; lys2-801; leu2-D1; his3-D200; trp1-D63; DYN1-3GFP::TRP;, ura3-52::CFP-TUB1::URA3; SPC110-tdTomato::SpHIS5; Flag-PAC1(E253A-H254A); ura3Δ::KanMX* | This work |
| RPY1797 | *MATa; lys2-801; leu2-D1; his3-D200; trp1-D63; DYN1-3GFP::TRP1; ura3-52::CFP-TUB1::URA3; SPC110-tdTomato::SpHIS5; Flag-PAC1(S248Q); ura3Δ::KanMX* | This work |
| RPY1798 | *MATa; lys2-801; leu2-D;1 his3-D200; trp1-D63; DYN1-3GFP::TRP;, ura3-52::CFP-TUB1::URA3; SPC110-tdTomato::SpHIS5; Flag-PAC1(F185D-I189D-R494A); ura3Δ::KanMX* | This work |
